# Glial control of sphingolipid levels sculpts diurnal remodeling of circadian circuits

**DOI:** 10.1101/2022.03.18.484007

**Authors:** John P. Vaughen, Emma Theisen, Irma Magaly Rivas-Sema, Andrew B. Berger, Prateek Kalakuntla, Ina Anreiter, Vera C. Mazurak, Tamy Portillo Rodriguez, Joshua D Mast, Tom Hartl, Ethan O. Perlstein, Richard J. Reimer, M. Thomas Clandinin, Thomas R. Clandinin

## Abstract

Structural plasticity in the brain often necessitates dramatic remodeling of neuronal processes and attendant reorganization of the cytoskeleton and membranes. While cytoskeletal restructuring has been studied extensively, how lipids might orchestrate structural plasticity remains unclear. We show that specific glial cells in *Drosophila* produce Glucocerebrosidase (GBA) to locally catabolize sphingolipids. Sphingolipid accumulation drives lysosomal dysfunction, causing *gba1b* mutants to harbor protein aggregates that cycle across circadian time and are regulated by neural activity, the circadian clock, and sleep. While the vast majority of membrane lipids are stable across the day, a specific subset, highly enriched in sphingolipids, cycles daily in a *gba1b*-dependant fashion. In parallel, circadian clock neurons remodel their neurites, growing and shrinking across the day to shape circadian behavior. Remarkably, this neuronal remodeling relies on a cycle of temporally offset sphingolipid biosynthesis and catabolism. Thus, dynamic sphingolipid regulation by glia enables diurnal circuit remodeling and proper circadian behavior.

## Introduction

Lifelong brain function requires coordinated biosynthetic and degradative pathways to precisely maintain neural membrane composition, circuit function, and animal behavior. While protein synthesis and degradation have been studied extensively in neurons, our understanding of lipid metabolism in the brain is comparatively limited. Brains are lipid-rich and display regional heterogeneity in lipid species (O’Brien and Sampson, 1965)(Fitzner et al., 2020), which include the CNS-enriched sphingolipids (Merrill, 2011). Moreover, neurons can dynamically reshape their membranes as part of structural and synaptic plasticity. For example, circadian pacemaker neurons in both vertebrates and *Drosophila* undergo dramatic daily cycles of membrane addition and removal (Krzeptowski et al., 2018)(Becquet et al., 2008)(Fernández et al., 2008) that correlate with cycles of behavioral activity (Petsakou et al., 2015) (Song et al, 2021). In addition, proteins that control membrane lipid composition are associated with Parkinson’s Disease and Alzheimer’s Disease (Yadav and Tiwari, 2014)(Lin et al., 2019) (Futerman and van Meer, 2004), and circadian dysfunction is a common feature of neurodegeneration that can precede disease onset (Leng et al., 2019). However, surprisingly little is known about how specific lipid species are coupled to structural plasticity in neuronal membranes.

A prominent class of neural membrane lipids are the glycosphingolipids, which are built upon a core of Glucosylceramide (GlcCer) from the sphingolipid Ceramide (Cer). GlcCer is degraded by Glucocerebrosidase (GBA) (Figure 1A), and mutations in *GBA* are linked to sleep disruption and neurodegeneration. Complete loss of GBA activity invariably causes neuropathic Gaucher disease, and *GBA* carriers are ~3-5 times more likely to develop Parkinson’s disease (Sidransky and Lopez, 2012) characterized by accelerated cognitive decline (Sidransky et al., 2009) (Liu et al., 2016). Mutations in *GBA* are also associated with isolated REM sleep behavior disorder (Gan-Or et al., 2015)(Krohn et al., 2020). However, the cellular mechanisms underpinning how mutations in *GBA* cause brain defects are unclear.

**Figure 1.**
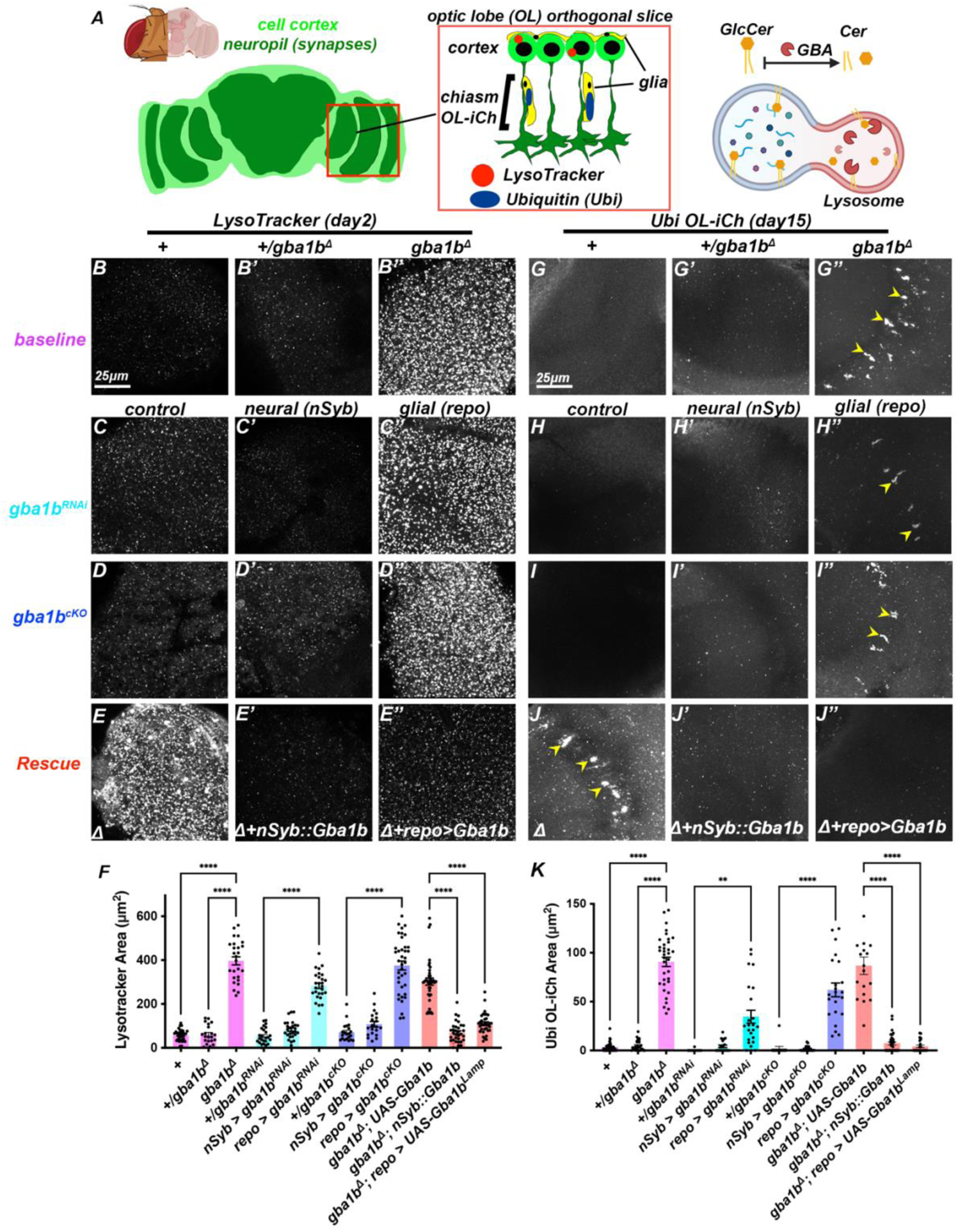
Gba1b is required in glia for neuronal lysosome function. (A) The *Drosophila* brain consists of neuronal and glial cell bodies arranged in a cortical rind (light green) enclosing dense synaptic neuropils (dark green). Within the Optic Lobe chiasm (OL-iCH), neuronal processes (dark green) are tightly associated with glial cells (yellow). Lysosomes (LysoTracker, red) are localized to cortical cell bodies (see *Figure S1*), while Ubiquitin (Ubi) aggregates are predominantly inside glia in *gba1b* mutant optic lobes (see *Figure S2*). Inside lysosomes, Gba1b degrades the lipid GlcCer into Cer. (B-E’’) LysoTracker labeling in brains from 2 day-old flies. (G-J’’) Ubiquitin aggregates in the OL-iCh in 15-20 day-old flies. (B, G) Control. (B’, G’) *gba1b^Δ1^*/+ heterozygotes. (B’’, G’’) *gba1b^Δ^* transheterozygotes have enlarged lysosomes and accumulate Ubi aggregates in the OL-iCh (arrows). (C-C’’, H-H’’) RNA-interference, *gba1b^RNAi^*. Control (C, H), neural (C’, H’; *nSyb-GAL4*), and glial (C’’, H’’; *repo-GAL4*). (D-D’’, I-I’’) Somatic *CRISPR*, *gba1b^cKO^*. Control (D, I), neural (D’, I’; *nSyb-GAL4*), and glial (D’’, I’’; *repo-GAL4*). Both *gba1b^RNAi^* and *gba1b^cKO^* in glial cells but not neurons cause enlarged Lysosomes and Ubi aggregates. (E-E’’, J-J’’) Null *gba1b^Δ^* (E, J) can be rescued by *nSyb::Gba1b* (E’, J’; neurons) or by overexpressing *UAS-Gba1b^LAMP^* in glia (E’’, J’’; *repo-GAL4*). (F) Quantification of LysoTracker data. (K) Quantification of Ubi data. *p<0.05, ****p<0.0001, ANOVA, Tukey’s multiple comparisons for normally distributed data, Kruskal-Wallis test for nonparametric data (E-E’’, G-J). n>20 optic lobes. Data are represented as mean ± SEM. Scale bar: 25μm. See also *Figure S1* and *Figure S2*, and *Table S1* for genotypes.

*Drosophila* is a powerful model for understanding GBA function: mutations in fly *gba1b* have evolutionarily conserved phenotypes, including lysosomal impairment, insoluble ubiquitin (Ubi) protein aggregates, and reduced sleep (Davis et al., 2016)(Kinghorn et al., 2016)(Kawasaki et al., 2017). Moreover, core sphingolipid regulatory enzymes (centered around Cer) are conserved in *Drosophila*, and mutations in these genes can cause neurodegenerative and behavioral changes (Acharya and Acharya, 2005) (Dasgupta et al., 2009)(Acharya et al., 2003). While prior studies identified a role for Gba1b in circulating extracellular vesicles (Jewett et al., 2021) as well as arthropod-specific glycosphingolipids in nervous system development (Haines and Irvine, 2005)(Huang et al., 2018)(Huang et al., 2016)(Soller et al., 2006)(Chen et al., 2007), the cellular requirements for the conserved breakdown of GlcCer by GBA in the adult brain remain unknown, and evidence for how sphingolipids sculpt neural circuitry and behavior is lacking *in vivo*.

We discover that *Drosophila* glia produce Gba1b to control lysosome homeostasis and sphingolipid degradation in adult neurons. Removal of *gba1b* from specific glial types triggered Ubi aggregate formation in both neurons and glia. Glia in both flies and humans have broadly conserved functions in neurotransmitter recycling, metabolism, and membrane phagocytosis (Freeman, 2015)(Yildirim et al., 2019). We identify neuropil ensheathing glia (EG), a glial type that is closely associated with neurites, as well as Perineural glia (PNG) as key mediators of sphingolipid catabolism. Remarkably, Ubi aggregates in young *gba1b* mutants shrink during the day and grow larger at night. This cycle of aggregate burden was modulated by the circadian clock and neural activity, as disrupting the circadian clock, or sleep perturbations, blocked aggregate cycling across the brain, and dark-rearing suppressed aggregate formation in the visual system. Comparative lipidomics studies at dusk and in the night revealed that while the levels of most membrane phospholipids were stable across the day, specific sphingolipid species fluctuated diurnally. Moreover, the cycles of neurite growth and retraction in the sLNv circadian pacemaker neurons strongly depended on sphingolipid regulation. Mutations in *gba1b,* or removal of *gba1b* from glia, blocked this remodeling and prevented neurite growth. Conversely, inhibiting sphingolipid biosynthetic enzymes selectively in sLNv cells abolished neurite retraction. Strikingly, reprogramming the temporal pattern of Gba1b by imposing elevated expression of Gba1b at night in these neurons was sufficient to invert the normal cycle of neurite remodeling, with increased terminal volume at dusk instead of dawn. Thus, glia degradation and neuronal biosynthesis of specific sphingolipids is both necessary and sufficient to sculpt axon terminals across the day.

## Results

### Gba1b is required in glia for neuronal lysosome function

To better characterize brain sphingolipid catabolism and Gba1b function, we generated an early frameshift allele, *gba1b^Δ1^*, which we used in *trans* to *gba1b^ΔTT^*, a characterized null allele that also deletes an adjacent gene, *Qsox4* (Figure S1A, Table S1, (Davis et al., 2016)). We used transheterozygous combinations of these alleles to produce viable adults designated *gba1b*^Δ^ (which retain one functional copy of Qsox4 and do not homozygous other chromosomal regions to avoid unwanted off-target effects from other loci). Consistent with previous work (Kinghorn et al., 2016), we confirmed that *gba1b*^Δ^ brains harbor enlarged degradative lysosomes marked by an acidic-compartment label (LysoTracker), active proteases (Cathepsin B), and a lysosomal protein (Lamp) (Figure 1B-B’’, Figure S1E-J). Aberrant lysosomes distributed across the cortex of the brain and were enclosed by neural but not glial membranes (Figure S1K-L). We exploited this robust neuronal phenotype to identify where Gba1b is required for lysosome maintenance. Unexpectedly, *gba1b* knockdown (*RNAi*) or knockout (somatic *CRISPR, gba1b^cKO^*) in all neurons did not cause any discernible lysosomal phenotypes (Figure 1C’, D’). In contrast, pan-glial *gba1b* knockdown or knockout triggered lysosomal hypertrophy (Figure 1C’’, D’’). These data argue that neuronal lysosome function depends on Gba1b expression in glia.

We next attempted to rescue the *gba1b* LysoTracker phenotype using three Gba1b over-expression constructs (Table S2, Figure S1A): a wild-type *Gba1b* transgene, a catalytically-inactive transgene (*Gba1b^E340K^*), and a tethered *Gba1b^Lamp^* transgene designed to restrict Gba1b to the lysosome (Mikulka et al., 2020). Glial overexpression of *Gba1b^Lamp^*or wild-type *Gba1b* rescued the LysoTracker phenotype (Figure 1E’’, Figure S1U). Importantly, expression of Gba1b in glia rescued the ectopic LysoTracker staining in neurons of *gba1b* mutants (Figure S1M-P), demonstrating that glial Gba1b is nonautonomously required to maintain neuronal lysosomes. In contrast, overexpressing Gba1b in neurons caused lysosomal hypertrophy in wild-type animals, a phenotype not seen upon overexpressing *Gba1b^E340K^* (Figure S1Q-R). Thus, excess Gba1b activity in neurons is detrimental to lysosomal homeostasis. To temper Gba1b induction in neurons, we placed Gba1b under the direct control of a neuronal enhancer (*nSyb::Gba1b*; Figure S1B-D). Interestingly, *nSyb::Gba1b* fully rescued the lysosomal phenotypes of *gba1b^Δ^* mutants (Figure 1E’). Thus, while Gba1b is selectively produced by glia and sufficient to rescue lysosomes in *gba1b*^Δ^ mutants, neurons can also utilize Gba1b.

We next tested if Gba1b was required in glia for other *gba1b^Δ^* phenotypes. A hallmark of *gba1b*^Δ^ brains is progressive accumulation of insoluble, polyubiquitinated proteins (Kinghorn et al., 2016)(Davis et al., 2016). We found that older *gba1b^Δ^* mutants accumulated Ubiquitin (Ubi) aggregates most prominently in both the inner optic lobe chiasm (OL-iCh) and in the cortex of the Mushroom Body calyx (MB-ca) (Figure 1G, Figure S2A-J), as well as in the blood-brain barrier (data not shown). Ubi extensively colocalized with Ubi-lysosomal adaptor protein p62, which is a common component of Ubi aggregates in fly and human neurodegeneration models (Bartlett et al., 2011) (Figure S2E-F). By expressing *p62-GFP* within neurons or glia in *gba1b^Δ^*brains, we determined that MB-ca puncta were neural, whereas larger Ubi aggregates were glial, including OL-iCh aggregates (Figure S2L, S2M). Similar to the LysoTracker phenotype, glial knockdown or knockout of *gba1b* induced Ubi aggregates, while neuronal perturbations did not (Figure 1H 1I, Figure S2J). Conversely, glial expression of *Gba1b^Lamp^* in *gba1b^Δ^* mutants fully rescued both Ubi aggregate populations (Figure 1J and Figure S1V, S1W). *nSyb::Gba1b* also rescued neuronal MB-ca aggregates (Figure S1W) and surprisingly also rescued glial OL-iCh aggregates (Figure 1J’). Taken together, Gba1b expression is required in glia to prevent lysosome dysfunction in neurons and subsequent Ubi aggregate appearance in both neurons and glia. However, Gba1b expression in neurons suffices to rescue *gba1b^Δ^*aggregate formation in both neurons and glia in the cortex.

### Specific glial subtypes are necessary and sufficient for Gba1b function in the brain

To identify which cells express Gba1b, we queried single-cell RNA-sequencing datasets and found that Gba1b transcripts were produced by multiple glial clusters during pupal development (Kurmangaliyev et al., 2020) (Figure S3A-A’). An adult dataset also revealed low levels of expression of Gba1b in glia but not neurons (Davie et al., 2018). To identify which glia cells express Gba1b, we characterized *Gba1b^GAL4^*, a gene trap in the Gba1b locus (Lee et al., 2018)*. Gba1b^GAL4^* drove expression of a fluorescent reporter in glia but never in neurons (Figure S3B-C). Combining *Gba1b^GAL4^* with a panel of *LexA*-based markers of individual glial types (Pfeiffer et al., 2008)(Kremer et al., 2017) revealed expression of Gba1b in multiple glial lineages, but not astrocyte-like glia (Figure S3D-K). We next removed Gba1b from individual glial subtypes using both RNAi and somatic CRISPR but found no LysoTracker phenotypes (Figure 2A, Figure S3L-M). However, given that Gba1b is expressed in multiple glia types, we reasoned that Gba1b may be redundantly required. We therefore screened combinations of glial cell *gba1b* knockouts and found that removing Gba1b from both EG and PNG caused lysosomal enlargement (Figure 2A’’). *gba1b^cKO^* in EG and PNG also caused Ubi aggregate accumulation in the OL-iCh and the MB-ca (Figure 2B-C). Conversely, expression of wild-type *Gba1b,* but not catalytically inactive *Gba1b^E340K^*, in EG was sufficient to rescue the lysosomal and Ubi aggregate phenotypes seen in *gba1b^Δ^* mutants (Figure 2D-F). However, expression of wild-type Gba1b in PNG failed to rescue these phenotypes (Figure 2D’’’-F’’’), and expression of Gba1b in astrocytes (which extensively tile the synaptic neuropil) failed to rescue LysoTracker (Figure 2G’). Importantly, EG, PNG, and astrocyte drivers expressed high levels of Gba1b protein in these rescue experiments (Figure S3N-P). Consistent with a critical role for Gba1b in EG, labeling EG membranes in *gba1b^Δ^* mutants revealed that Ubi aggregates in the OL-iCh and MB-ca were closely apposed to EG cells (Figure 2H-I). Thus, Gba1b is required in EG and PNG, and expression in EG is sufficient to rescue neuropil and cortex phenotypes.

**Figure 2.**
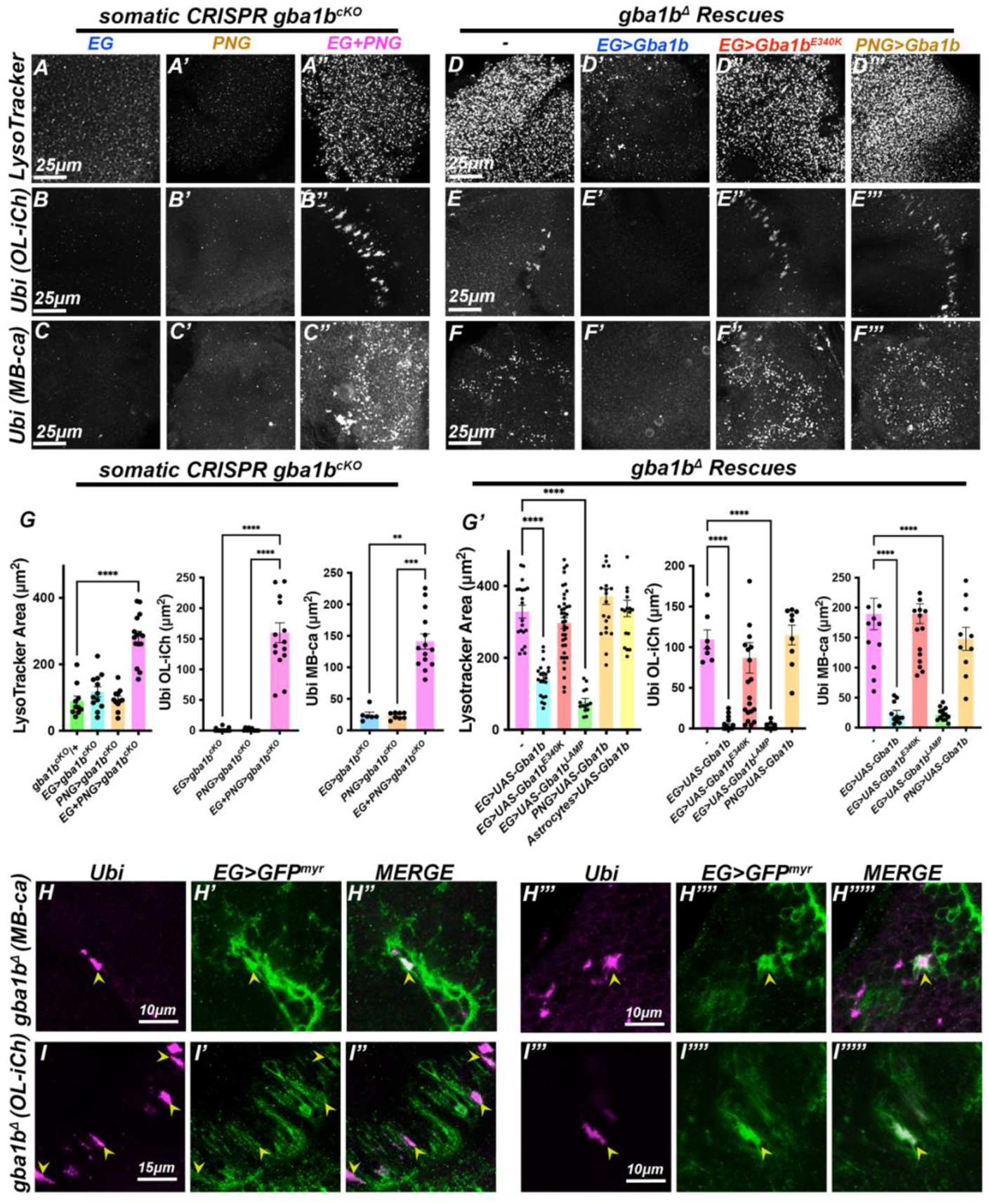
Specific glial subtypes are necessary and sufficient for Gba1b function in the brain. (A-C’’) *gba1b* somatic *CRISPR* (*gba1b^cKO^)* in ensheathing glia (EG), perineural glia (PNG), or both EG and PNG (see Figure S3D for anatomy diagram). (D-F’’’) Rescue of *gba1b^Δ^* by overexpressing Gba1b in EG and PNG. (A-A’’, D-D’’’) LysoTracker labeling in 2 day-old brains. (B-F’’’) Ubi labeling in 20-25 day-old brains. (A, B, C) *gba1b^cKO^*in EG. (A’, B’, C’) *gba1b^cKO^* in PNG. (A’’, B’’, C’’) *gba1b^cKO^*in both EG and PNG, a perturbation that caused enlarged lysosomes and Ubi aggregates. (D’, E’, F’) *gba1b^Δ^* rescue by wild-type *UAS-Gba1b* expression in EG but not by enzyme-dead *UAS-Gba1b^E340K^* (D’’, E’’, F’’). (D’’’, E’’’, F’’’) PNG expression of wild-type *UAS-Gba1b* failed to rescue *gba1b^Δ^*. (G) Quantification of LysoTracker data, including the control ISOD1/*gba1b^cKO^*(green), as well as Ubi in the Ol-ich and MB-ca. (G’) Quantification of rescue data (including data from *EG>Gba1b^LAMP^* (green) and Astrocytes overexpressing Gba1b (yellow) for LysoTracker). (H-H’’’’’) EG membranes (green) and subsets of Ubi (magenta) in *gba1b^Δ^* colocalize in the MB-ca (arrowheads). (I-I’’) EG membranes (green) enclose Ubi (magenta) in *gba1b^Δ^* OL-iCh. (I’’’-I’’’’’) Subsets of OL-iCh Ubi structures accumulate bright EG membrane in *gba1b^Δ^* (arrowheads). *p<0.05, ****p<0.0001, ANOVA, Tukey’s multiple comparisons for normally distributed data, Kruskal-Wallis test for nonparametric data (B-C’’, E-F’’’). n>6 brains. Scale bar: 25μm (A-F); 10μm (H-H’’’’’, I’’’-I’’’’’), 15μm (I-I’’). Data are represented as mean ± SEM. See also *Figure S3*, and *Table S1* for genotypes.

### Ubiquitin aggregates cyclically grow and shrink at younger ages

We next examined the onset of Ubi aggregate deposition in young *gba1b^Δ^* flies and found an unanticipated relationship between aggregate burden and circadian time. Specifically, Ubi aggregates in the OL-iCh and the MB-ca were low or absent during the day but accumulated at night in 7-day old *gba1b* mutants (Figure 3A-F). We more closely examined the relationship between aggregates, age, and circadian zeitgeber time (ZT). For both *gba1b^Δ^* and pan-glial *gba1b^cKO^* animals, the nadir of aggregate burden occurred at ZT6 (afternoon), and aggregates grew steadily through ZT12 (evening) and ZT18 (night) in the OL-iCh (Figure 3G-L). Similarly, Ubi aggregates in the MB-ca were smaller during the day and larger at night in young flies (Figure S4A-D). However, in older flies, the difference between these timepoints was less pronounced (Figure 3H, K and Figure S4C), arguing that diurnal aggregate clearance diminishes with age. In contrast to Ubi aggregates, lysosome markers were not as dramatically modulated by the circadian clock in *gba1b^Δ^* null animals (Figure S4E-G). Subtle changes in control lysosome morphology were detected across time in controls, consistent with circadian autophagosome production (Bedont et al., 2021)(Ryzhikov et al., 2019) (Ulgherait et al., 2021) (Figure S4H-J). Thus, Ubi aggregates but not lysosomes undergo significant variations in size in *gba1b* mutants that tracked with circadian time.

**Figure 3.**
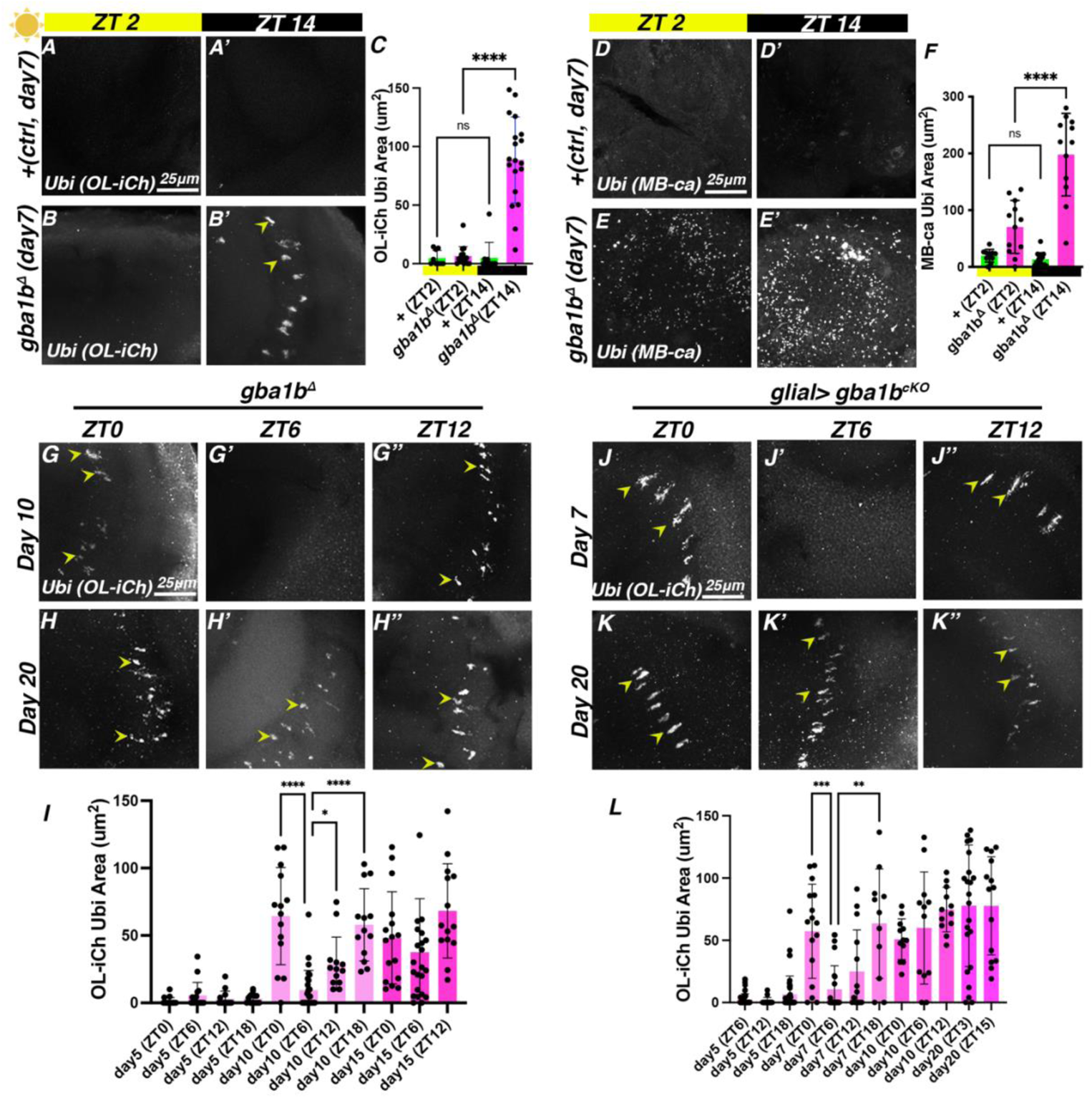
Ubiquitin aggregates cyclically grow and shrink at younger ages. Ubiquitin labeling (Ubi, white) in controls (A, A’, D, D’) and *gba1b^Δ^* (B, B’, E, E’) examined in both the OL-iCh and the Mushroom Body calyx (MB-ca) at zeitgeber time ZT2 (day) and ZT14 (night). (C) Quantification of A-B’. (F) Quantification of D-E’. (G-H) Ol-iCh Ubi in *gba1b^Δ^* and (I-J) glial *gba1b^cKO^* examined at ZT0, ZT6, and ZT12 at younger (G, J) and older (H, K) ages. (I) Quantification of Ubi aggregates in *gba1b^Δ^* across days and ZT. (L) Quantification of Ubi aggregates in glial *gba1b^cKO^*across days and ZT. Younger *gba1b^Δ^*and glial *gba1b^cKO^* brains accumulate Ubi at night and have a midday nadir (ZT6), whereas older *gba1b* flies plateau aggregates at an elevated level. *p<0.05, ****p<0.0001, Kruskal-Wallis test for nonparametric data, multiple comparisons. n>15 brains. Scale bar: 25μm. Data are represented as mean ± SEM. See also *Figure S4*, and *Table S1* for genotypes.

### Ubiquitin aggregate size is controlled by neural activity and the circadian clock

We hypothesized that the aggregate cycle apparent in *gba1b^Δ^* flies could be directly controlled by the circadian clock, by light-evoked changes in neural activity (which entrains the clock), or by both. We tested these possibilities by depriving flies of light and by mutating *period* (*per*), a core component of the circadian clock (Figure 4A). Dark-rearing *gba1b^Δ^* flies (DD) dramatically suppressed the number of Ubi aggregates in the OL-iCh, even in 20 day-old flies (Figure 4B, D). In contrast, Ubi aggregates in the MB-ca persisted in DD conditions (Figure 4C, E). This suggests that Ubi aggregates in the OL-iCh are sensitive to light and subsequent neural activity. In contrast, Ubi aggregates in the MB-ca are not responsive to light deprivation, perhaps because light does not strongly modulate neural activity in this region.

**Figure 4.**
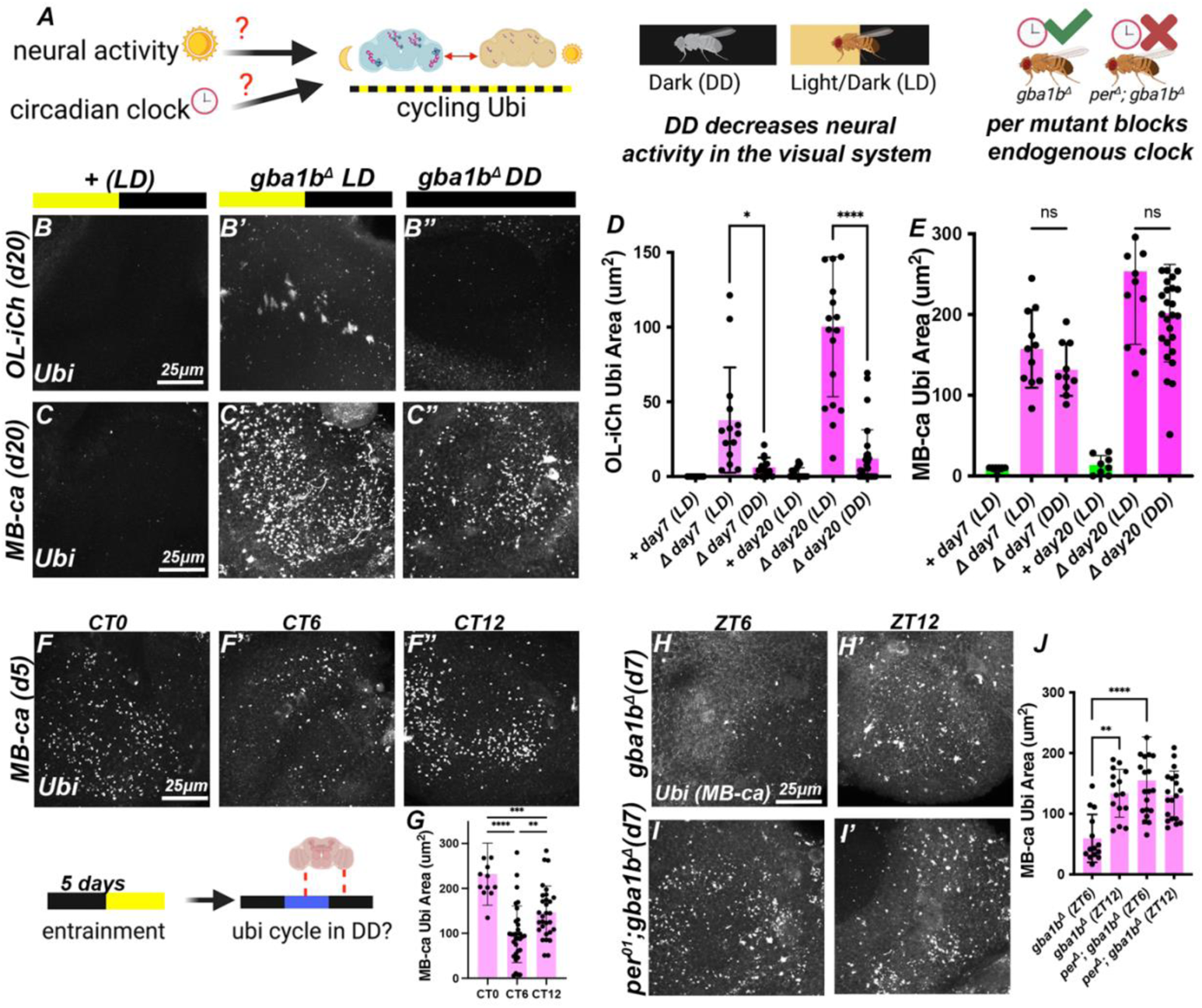
Ubiquitin aggregate size is controlled by neural activity and the circadian clock. (A) Schematic of hypothesis that cycling Ubi aggregates in young *gba1b^Δ^*flies might be controlled by light and/or the circadian clock. (B-E) Dark-rearing reduces Ubi aggregate formation in the Ol-iCh (B) but not the MB-ca (C). (B,C) Control. (B’,C’) *gba1b^Δ^* raised under light-dark conditions (LD). (B’’, C’’) *gba1b^Δ^* raised under dark-dark conditions (DD). (D) Quantification of Ubi in the OL-iCh. (E) Quantification of Ubi in the MB-ca. Dark-rearing suppressed Ubi aggregate formation in the OL-iCh but not the MB-ca in *gba1b^Δ^*. (F) Ubi labeling (white) in *gba1b^Δ^* at three different circadian times (CT0, CT6, CT12) in the MB-ca after shifting entrained flies to DD conditions. (G) Quantification of MB-ca Ubi burden reveals nadir at CT6. (H-I) Ubi labeling (white) in *per^01^; gba1b^Δ^* double mutants (I) at ZT6 and ZT12 compared to *gba1b^Δ^* single mutants. We observed that the midday nadir of Ubi aggregate formation was lost in *per^01^; gba1b^Δ^* double mutant animals compared to controls (quantified in J). n>15 brains. *p<0.05, ****p<0.0001, ANOVA, Tukey’s multiple comparisons for normally distributed data, and Kruskal-Wallis test for nonparametric data (B-E). Scale bar: 25μm. Data are represented as mean ± SEM. See also *Figure S4*, and *Table S1* for genotypes.

To test whether aggregate formation is under direct control of the endogenous circadian clock without the confound of light sensitivity, we focused on aggregate formation in the MB-ca. We entrained flies in LD, shifted to constant darkness for one day, and still detected lower aggregate burden at circadian time CT6 (Figure 4F-G). Moreover, introducing the arrhythmic *per*^01^ allele (Konopka and Benzer, 1971) into the *gba1b^Δ^* background abolished cyclic changes in Ubi aggregate burden (Figure 4H-J). While mutating the circadian clock alone did not cause Ubi aggregates to accumulate in the OL-iCh (Figure S4L), we found enhanced Ubi burden in PNG and OL-iCh glia in *per*^01^*; gba1b^Δ^* flies (Figure S4K). Consistent with brain activity regulating aggregate deposition, sleep-depriving *gba1b^Δ^* flies at night plateaued Ubi cycling in both MB-Ca and OL-iCh (Figure S4M-N), similar to the effects of clock disruption via *per^01^*. Moreover, short-sleeping mutants *sleepless* and *insomniac* (Koh et al., 2008)(Stavropoulos and Young, 2011) also progressively accumulated Ubi aggregates in the MB-ca (Figure S4O-P), further suggesting that altered neural activity following sleep disruptions influences proteostasis. In sum, *gba1b^Δ^* neuronal aggregate size in the MB-ca is directly controlled by the circadian clock, and glial aggregate size in the OL-iCh is controlled by both neural activity and the circadian clock.

### GBA regulates specific sphingolipids and phospholipids

As Gba1b degrades the lipid GlcCer, we tested which brain lipids were differentially controlled by Gba1b. We first examined an antibody directed against GlcCer alongside neutral lipids using the lipophilic Nile Red stain. Both GlcCer and Nile Red were increased in *gba1b^Δ^*brains compared to control animals (Figure 5A, B), and GlcCer was increased in pan-glial and EG + PNG *gba1b^cKO^* brains (Fig S5A). Ectopic Nile Red-positive lipids in *gba1b^Δ^* co-localized with LysoTracker (Figure 5C) and also colocalized with GlcCer (Figure 5D). Thus, GlcCer accumulation in *gba1b^Δ^* mutants localizes to engorged, lipid-filled lysosomes.

**Figure 5.**
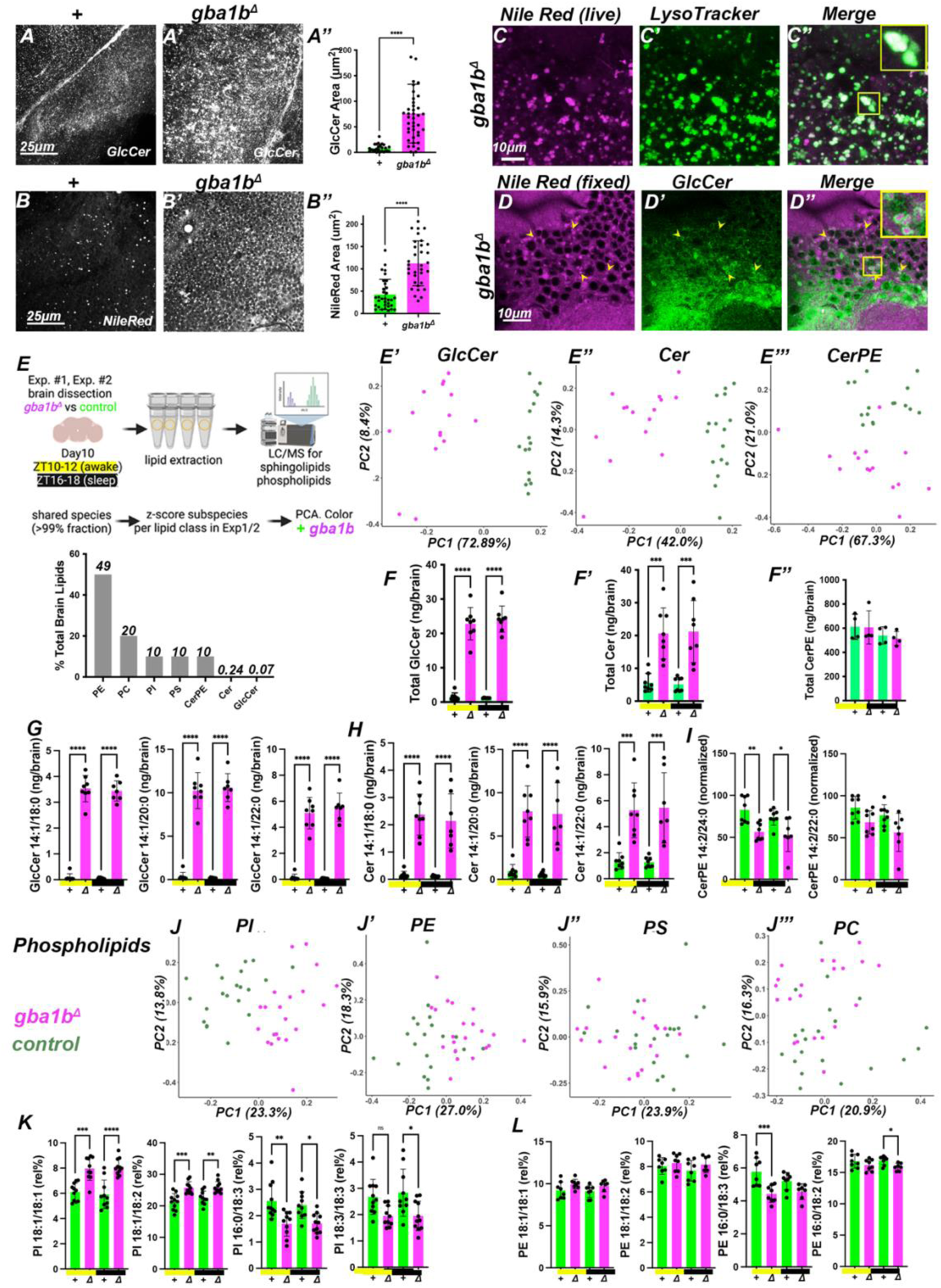
Gba1b regulates specific sphingolipids and phospholipids. (A) Staining against GlcCer in OL cortex in control (A) and *gba1b^Δ^* (A’) at ZT12 in 5 day-old brains. (A’’) Quantification of GlcCer area. *gba1b^Δ^* mutants have increased GlcCer. (B) Staining against neutral lipids with Nile Red in control (B) and *gba1b^Δ^* (B’) at ZT12 in 5 day-old brains. (B’’) Quantification of Nile Red. *gba1b^Δ^* mutants have increased neutral lipid burden. (C) Co-labeling *gba1b^Δ^* mutants with Nile Red (magenta) and LysoTracker (green) in the OL. LysoTracker particles are Nile Red positive (inset). (D) Fixed brains co-stained for Nile Red (magenta) and GlcCer (green) revealed that GlcCer encircled Nile Red puncta (arrowheads and inset). (E) Lipidomic analysis of fly brains from control and *gba1b^Δ^* animals, dissected in day 10 old flies at two timepoints, ZT10-12 (awake, peak evening activity) and ZT18-20 (midnight). Sphingolipids and phospholipids were extracted from two independent experiments and analyzed by liquid chromatography mass spectroscopy (LC-MS). Relative % fraction estimates of analyzed lipids are shown in the bar plot (bottom), with PE and PC dominating and Cer and GlCer exceedingly scarce. (E-E’’) Principal Component Analysis (PCA) of sphingolipids z-scored by abundance per sphingolipid species and colored by genotype (green = control, magenta = *gba1b^Δ^*). (E’) GlcCer biplot of PC1 and PC2 separates samples by genotype. (E’’) Cer separated *gba1b^Δ^* samples from controls. (E’’’) CerPE separated *gba1b^Δ^* samples from controls but more weakly. (F-F’’) Total levels of sphingolipid classes (ng/brain) revealed large GlcCer and Cer accumulation but no changes to bulk CerPE at day 10 for both timepoints (yellow bar denotes ZT10-12, dark bar denotes ZT16-18). (G-H) Specific species of GlcCer (G) and Cer (H) that accumulate in *gba1b* mutants and disproportionately contribute to the total GlcCer/Cer burden. (I) Some species of CerPE are modestly decreased in *gba1b^Δ^* mutants. (J-J’’’) PCA of phospholipids, z-scored by abundance per species and colored by genotype (green = control, magenta = *gba1b^Δ^*). (J) PI clearly separated samples by genotype, while PE (J’) modestly did so as well. PC (J’’) and PS (J’’’) did not separate *gba1b^Δ^* samples from controls. (K) Bidirectional regulation of PI species by *gba1b^Δ^.* (L) The same PE species were also reciprocally modulated by *gba1b^Δ^*. *p<0.05, ****p<0.0001, ANOVA, Tukey’s multiple comparisons for normally distributed data. n>10 (A-D) and n=15 brains/point, 8 biological replicates (E-I). Data are represented as mean ± SEM. See also *Table S1* for genotypes, and *Table S3* for raw lipidomics data.

We next conducted extensive lipidomics of *gba1b^Δ^* and control brains analyzed using liquid chromatography/mass spectrometry to detect sphingolipids (GlcCer, Cer and Ceramide-phosphatidylethanolamine (CerPE)) and phospholipids (Phosphatidylinositol (PI), Phosphatidylethanolamine (PE), Phosphatidylcholine (PC), and Phosphatidylserine (PS); Figure 5E). Given that we detected circadian aggregates and that subsets of lipids can vary across circadian time in non-neuronal cells(Katewa et al., 2016), we dissected brains from two timepoints, ZT10-12 (peak of evening activity) and ZT16-18 (midnight). Sphingolipids and phospholipids are classified by carbon chain length (for each chain) and the number of carbon-carbon double bonds. We detected 163 sphingolipid and phospholipid species (Table S3) which we divided into classes (GlcCer, Cer, CerPE, PI, PS, PC, and PE) and then analyzed by Principal Component Analysis (PCA).

PCA revealed very large differences in GlcCer and Cer between control and *gba1b^Δ^* mutant animals (73% of variance in GlcCer, 42% of variance in Cer; Figure 5E-E’). CerPE also modestly differentiated samples by genotype, accounting for 20% of variance (Figure 5E’’’). Consistent with the role of GBA in sphingolipid catabolism, total GlcCer and Cer levels were markedly increased in *gba1b^Δ^* mutant brains as well as pan-glial *gba1b^cKO^* brains (Figure 5F-F’’, Figure S5B). These changes were largely driven by three GlcCer and Cer species with identical carbon chains: 14:1/18:0, 14:1/20:0, and 14:1/22:0 (Figure 5G-H), representing ~80% total GlcCer and ~70% total Cer in *gba1b* mutants. In contrast, CerPE displayed little change in *gba1b^Δ^* mutants, with modest decreases in a handful of species (Figure 5I).

We next analyzed phospholipids (Figure 5J). Although bulk phospholipid levels were unchanged in *gba1b^Δ^* mutants (Figure S5C), PI and PE separated samples by genotype, accounting for ~20% of variance along PC1 (Figure 5J-J’’). PC and PS did not differentiate samples by genotype in any PCs (Figure 5J’’-J’’’; data not shown). Given that bulk phospholipids were unchanged, yet PCA separated PI and PE by *gba1b^Δ^* and control samples, we reasoned that specific species of PE or PI might reciprocally change in *gba1b^Δ^* mutants. Indeed, in *gba1b^Δ^* brains, 18:1/18:1 and 18:1/18:2 PI and PE species were increased, whereas 16:0/18:3 PE and PI were decreased (Figure 5K-L). In total, *gba1b* mutants significantly modulated 27 GlcCer, 20 Cer, 11 PI, 6 CerPE, 4 PE, 3 PC, and 3 PS species (Table S4). Notably, the unchanged PS, PC, and PE species account for >65% of total brain lipids (Guan et al., 2013)). Thus, Gba1b exerts specific effects on subsets of brain lipids, predominantly GlcCer and Cer species, alongside selective effects on CerPE, PI, and PE species.

### Sphingolipid levels fluctuate diurnally

We next explored whether any lipids changed across the day by running PCA on sphingolipids and phospholipids from control animals (Figure 6A). Strikingly, the first two principal components for GlcCer and CerPE species strongly separated samples taken during the day and night, as did Cer, albeit more weakly (Figure 6B). In contrast, time of day accounted for little variance in any of the phospholipids (Figure 6C; data not shown).

**Figure 6:**
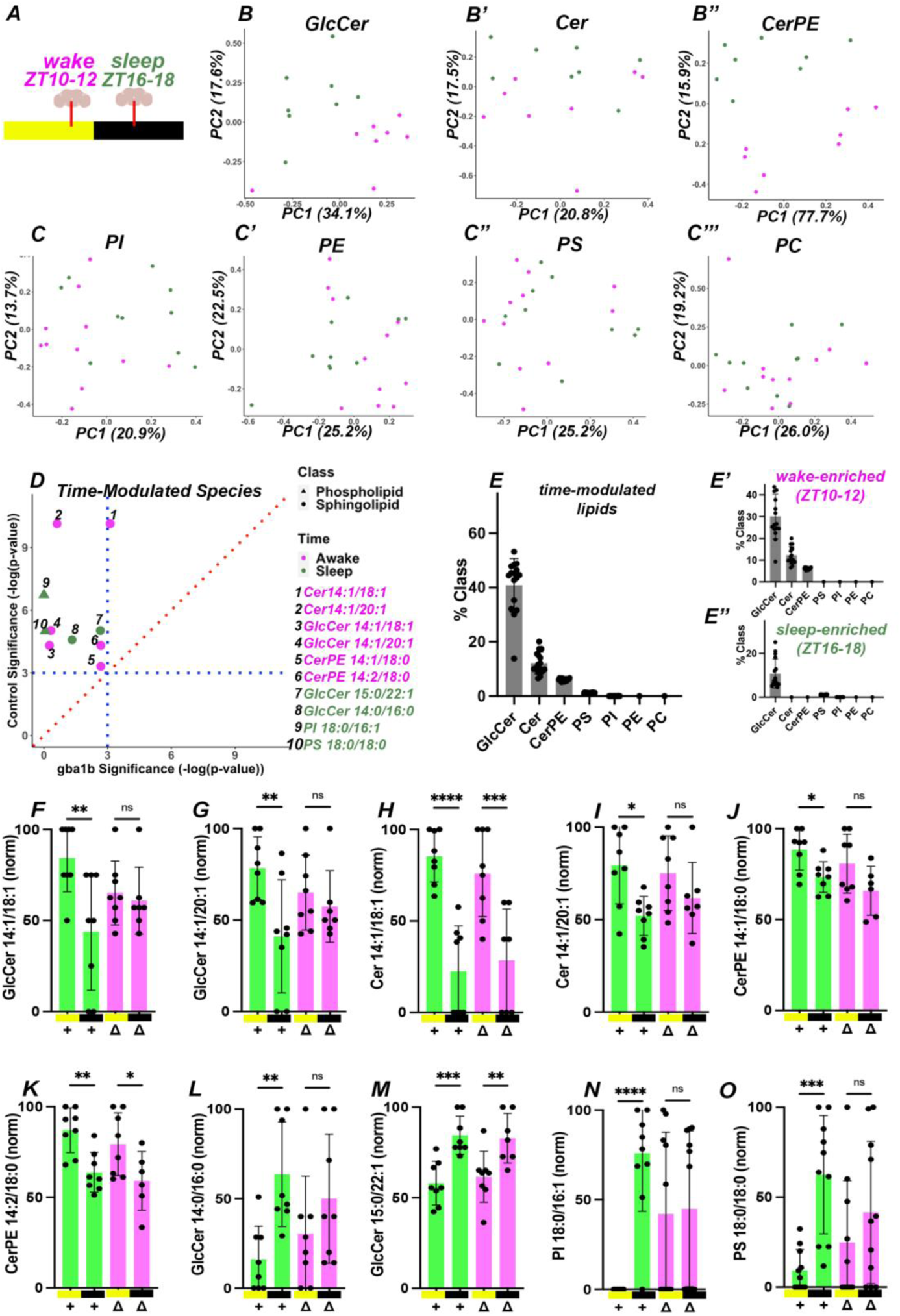
Sphingolipid levels fluctuate diurnally. (A) Principal Component Analysis (PCA) of sphingolipids and phospholipids z-scored by abundance per species in 10 day old control brains, colored by time (green = sleep/ZT16-18, magenta = wake/ZT10-12). GlcCer (B). Cer (B’). CerPE (B’’). GlcCer and CerPE strongly split samples by time, while Cer did so only modestly. (C-C’’’) Phospholipids. (C) PI. (C’) PE. (C’’) PS. (C’’’) PC. Phospholipids did not show strong circadian modulation. (D) Mean matrices for ZT16-18/night and ZT10-12/day samples were constructed and subtracted to identify circadian modulated species in controls with FDR <0.05 (dotted blue lines, see *STAR Methods*). Control species significance (y-axis) for diurnal fluctuation in these time-modulated species is plotted against significance in *gba1b^Δ^* samples (x-axis). Sphingolipid species are plotted as circles and phospholipids as triangles; wake-enriched species are colored green, sleep-enriched are colored magenta. In controls, ten species significantly fluctuate across time, eight of which were sphingolipids (also see *Table S4*). (E) Diurnally modulated species contributed to ~40% of total GlcCer, 10% of total Cer, and 6.5% of total CerPE. Diurnally modulated PS and PI contributed to 1% of PS and 0.4% PI, respectively. (E’-E’’) Data from (E) split into ‘awake-enriched’ and ‘sleep-enriched’. (F-O) Normalized values of the ten time-modulated species shown for controls (green) and *gba1b* (magenta), with *gba1b^Δ^* diminishing the diurnal fluctuation across all species. *p<0.05, ****p<0.0001, ANOVA, Tukey’s multiple comparisons for normally distributed data. n=15 brains/point, 8 biological replicates. Data are represented as mean ± SEM. See also *Table S1* for genotypes, and *Table S3* for raw lipidomics data.

To directly reveal which lipid species were time-regulated, we constructed mean matrices from day and night samples and identified the largest differences between these vectors (FDR<0.05; see STAR Methods). Consistent with the results from PCA, of the ten species with circadian differences in control animals, eight were sphingolipids (Figure 6D-O). All ten species displayed reduced diurnal modulation in *gba1b^Δ^* mutants (Figure 6D-O). Crucially, the eight modulated sphingolipids represented 40% of GlcCer, 10% of Cer, and 6.5% of CerPE species (Figure 6E-E’’). Moreover, two of the GlcCer and Cer species enriched during activity at dusk have the same fatty acyl carbon chain compositions (14:1/18:1 and 14:1/20:1), suggesting common functions, or coupled biosynthesis. In contrast, sleep-enriched species appeared to feature more saturated chains (green species, Figure 6D; and see Table S4). Finally, we note that time-modulated species differ from those most upregulated in *gba1b* mutants. For example, GlcCer 14:1/18:0 was upregulated by *gba1b^Δ^* (Figure 5G), whereas GlcCer 14:1/18:1 instead fails to fluctuate with time in *gba1b^Δ^*(Figure 6F). Thus, circadian rhythms selectively affect sphingolipids, particularly GlcCer, and this modulation is strongly suppressed in *gba1b* mutants.

### Sphingolipid catabolism is required for adult sleep behavior

As *gba1b^Δ^* mutants have altered lipid profiles with blunted circadian fluctuations in lipid species, we reasoned that *gba1b* mutants might display defects in circadian behaviors including activity and sleep. We tested whether *gba1b^Δ^* mutants had deficiencies in circadian rhythms or sleep by measuring daily activity patterns. Control flies are crepuscular, with two bouts of activity surrounding dawn and dusk, with both sexes sleeping more at night than during the day (Figure 7A, (Helfrich-Förster, 2000)). Like controls, *gba1b* mutants began sleeping shortly after lights-off, but both sexes displayed diminished sleep characterized by shorter sleep bout duration (Figure 7A, B; Figure S6A, B). Pan-glial *gba1b* knockouts also displayed sleep loss associated with reduced sleep bout number rather than length (Figure 7C, Figure S6C). Moreover, removing *gba1b* specifically in both EG and PNG also reduced sleep (Figure 7D, Figure S6D). Depleting *gba1b* only in EG did not cause obvious sleep defects, while depleting *gba1b* in PNG only reduced daytime sleep (Figure S6I), consistent with barrier glia regulating sleep behavior (Artiushin et al., 2018)(Kozlov et al., 2020). Importantly, expressing Gba1b in both EG and PNG rescued the sleep deficits seen in *gba1b^Δ^*mutants (Figure 7E, Figure S6E), whereas individual PNG or EG drivers had only modest effects (Figure S6J).

**Figure 7.**
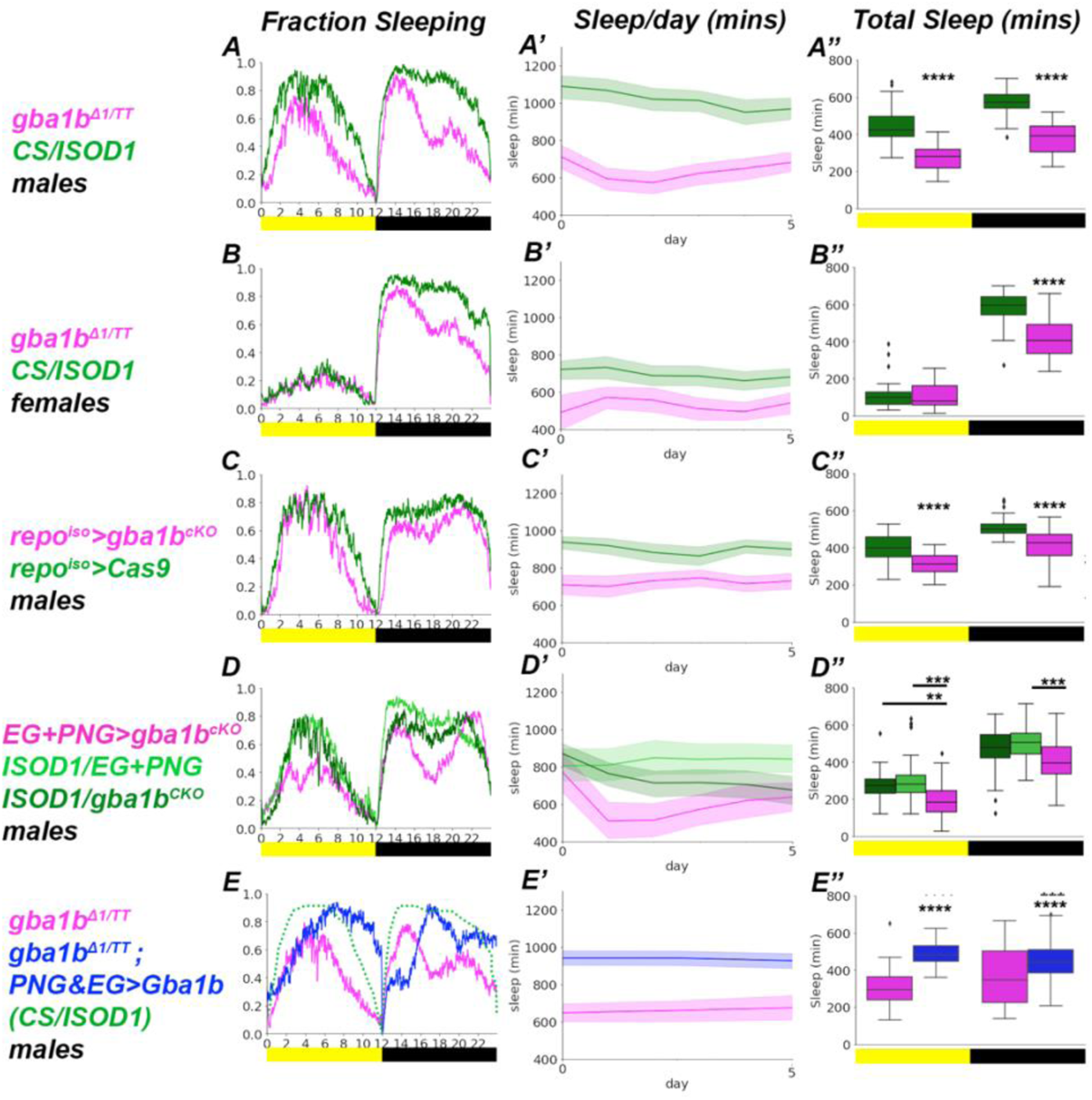
Sphingolipid catabolism is required for adult sleep behavior. *Drosophila* activity-monitor (DAM) assays for measuring sleep. Adult flies were analyzed from day 5 to day 10 on a 12-hour Light-Dark (LD) cycle. (A-A’’) Fraction of flies asleep across ZT for male *gba1b^Δ^* flies (magenta) versus control (CS/ISOD1) males (green). (B-B’’) Fraction of flies asleep across ZT for female *gba1b^Δ^* flies (magenta) versus control (CS/ISOD1) females (green). (C-C’’) Fraction of flies asleep across ZT for glial *gba1b^cKO^* males (magenta) versus control males (repo>cas9). (D-D’’) Fraction of flies asleep across ZT for EG+PNG *gba1b^cKO^* males (magenta) versus control males (ISOD1/EG+PNG, and ISOD1/ *gba1b^cKO^*). (E-E’’) Fraction of flies asleep across ZT for male *gba1b^Δ^* flies (magenta) versus rescue (EG+PNG>Gba1b) males (blue); ISOD1 control is overlaid (green, from A). (A, B, C, D, E) Fraction sleeping. (A’, B’, C’, D’, E’) Sleep per day. (A’’, B’’, C’’, D’’, E’’) Total Sleep. Reducing Gba1b activity via either *gba1b^Δ^*, or *repo-GAL4>gba1b^cKO^*, or EG+PNG *gba1b^cKO^* caused reduced sleep across days and times. EG+PNG expression of Gba1b was sufficient to partially rescue *gba1b^Δ^*, suggesting that there may be additional sleep deficits caused by metabolic functions of Gba1b outside the brain. *p<0.05, ****p<0.0001, ANOVA, Tukey’s multiple comparisons or Welch’s ANOVA depending on equality of data variance. For nonparametric sleep bouts, Mann-Whitney U test was used. n>28 flies. Data are represented as mean ± SEM, as well as the IQR for the Total Sleep boxplots. See also *Figure S6*, and *Table S1* for genotypes.

We next tested for persistence of circadian rhythms when *gba1b* mutants were moved to continuous darkness (DD) after entrainment in LD. Although we observed timely evening and morning peaks of activity (Figure S6F-G), arguing against complete abrogation of the circadian clock, *gba1b^Δ^* mutants and glial *gba1b* removal (*repo* or *EG+PNG)* had significantly weaker rhythmicity when free-running in darkness compared to controls (Table S5). Thus, loss of *gba1b* causes both sleep and circadian defects, and expression of Gba1b in the EG and PNG is necessary for proper rhythm strength as well as sleep behavior.

### Sphingolipids drive cyclic remodeling of sLNv neurites

We hypothesized that some of the behavioral deficits seen in *gba1b^Δ^* mutants might be linked to a failure to properly remodel circadian circuits. sLNv peptidergic neurons autonomously express pigment-dispersing factor (PDF) and undergo dramatic daily remodeling of neurites and terminals (Fernández et al., 2008) essential for the proper regulation of circadian patterns (Grima et al., 2004)(Parisky et al., 2008)(Choi et al., 2012) ((Valadas et al., 2018). In control animals, sLNv neurites are enlarged at dawn and smaller at dusk, a robust diurnal cycle (Figure 8A; (Petsakou et al., 2015)). However, in *gba1b^Δ^* mutants, sLNv neurites were trapped at an intermediate size in 5-day and 10-day old animals, with reduced sLNv volume and 3-D spread (Figure 8B, Figure S7A-B, Figure 8J). Critically, knocking-out *gba1b* in EG and PNG also trapped sLNv neurites at a constant reduced size (Figure 8C, Figure S7C). We next examined cell-type specific rescue of sLNv remodeling in *gba1b^Δ^* mutants. *nSyb::Gba1b,* which expresses Gba1b pan-neuronally, rescued sLNv remodeling in *gba1b^Δ^* mutants (Figure 8D). While expressing Gba1b in EG or PNG alone failed to rescue cyclic remodeling of sLNv terminals in *gba1b^Δ^* mutants (Figure S7D-E), expressing Gba1b in both EG and PNG significantly rescued axonal volume (Figure 8E) but not 3-D spread (Figure 8J-J’). This requirement for Gba1b in both EG and PNG is consistent with the close associated of sLNv neurites with both of these glial subtypes (data not shown). Thus, Gba1b is required non-autonomously in EG and PNG glia to remodel circadian circuitry, and can act directly in sLNv neurons, given rescue by ectopic expression using *nSyb::Gba1b*.

**Figure 8.**
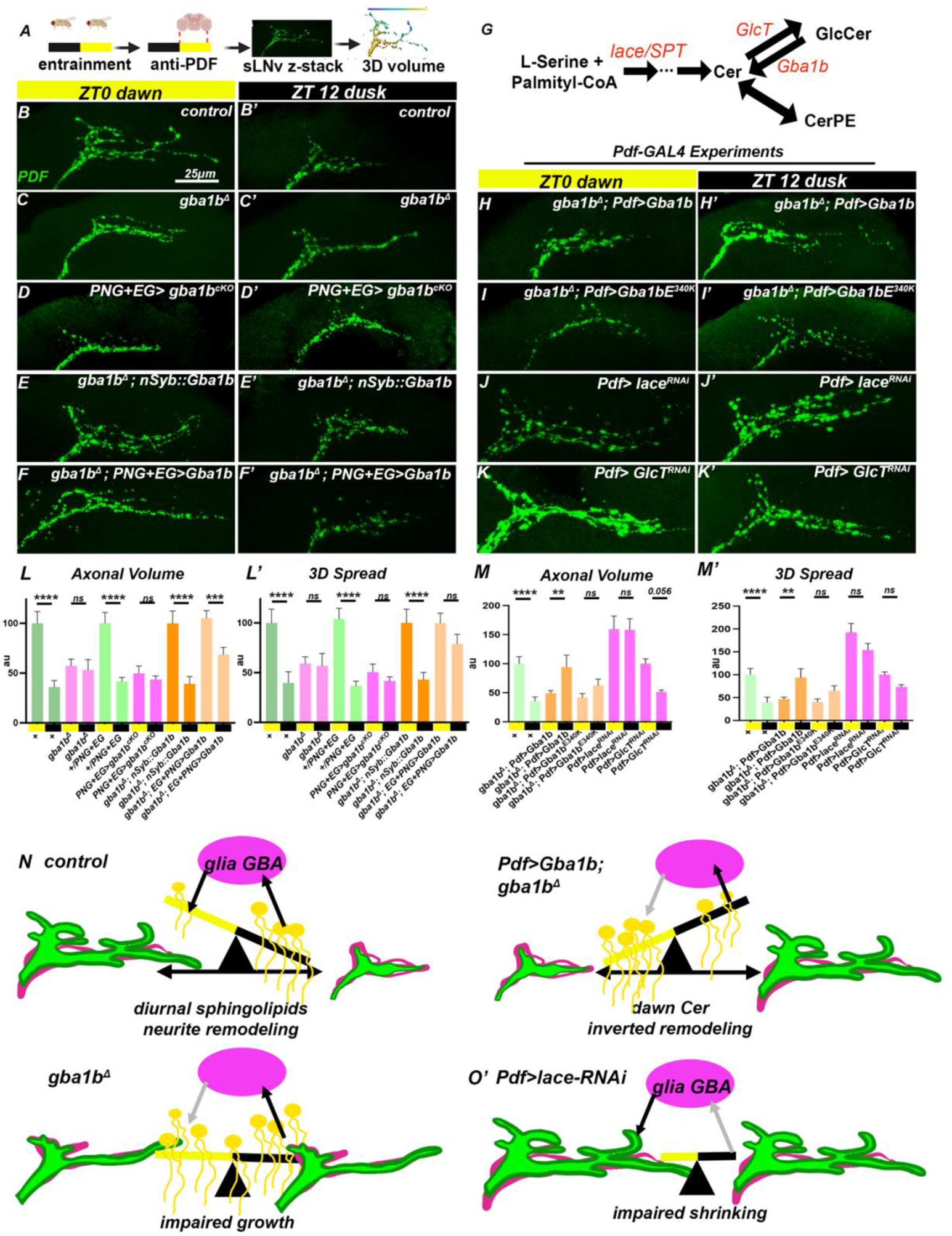
Sphingolipids drive cyclic remodeling of sLNv neurites. (A) Schematic of the assay for circadian sLNv neurite remodeling using anti-PDF, which faithfully represents sLNv neurite morphology (Fernández et al., 2008). Following circadian entrainment, control and *gba1b^Δ^* brains were dissected at ZT0 (8AM, dawn) and ZT12 (8PM, dusk) and stained with anti-PDF, before acquiring 3D volumes of neurite spread. (B) Confocal projections of sLNv neurites from 5-day old controls. Neurites are spread at dawn (ZT0) but retracted by dusk (ZT12). (C) *gba1b^Δ1^* mutants and (D) *PNG+EG>gba1b^cKO^* neurites fail to cycle sLNv neurites compared to controls. (E) *nSyb::gba1b* fully rescued sLNv remodeling defects of *gba1b^Δ^* mutants. (F) EG+PNG expression of Gba1b only rescued sLNv volume cycling in *gba1b^Δ^* mutants. (G) Simplified schematic of sphingolipid biosynthesis and degradation centered on Cer. (H) Expressing Gba1b in sLNv neurites in *gba1b^Δ^* using *Pdf-GAL4* inverted the remodeling cycle, with larger sLNv spread at dusk. (I) Expressing catalytically-dead *Gba1b^E340K^* with *Pdf-GAL4* in *gba1b^Δ^*did not cause inversion of sLNv cycling. (J) Impairing *de novo* sphingolipid biosynthesis in sLNv using *Pdf>lace^RNAi^* drastically increased neurite volume and blocked remodeling. (K) Impairing GlcCer biosynthesis in sLNv (*Pdf>GlcT^RNAi^*) modestly impaired cycling in neurite volume and blocked 3-D spread. (L-M) Quantification of sLNv neurites using 3D Spread and axonal volume. Controls from L are duplicated in M; experiments were normalized to highest-volume conditions run in parallel. (N) Model describing how diurnal changes in sphingolipid levels instruct sLNv neurite remodeling. In intact animals, sphingolipids (GlcCer/Cer) accumulate across the day to promote neurite retraction at dusk. EG/PNG glia express Gba1b (magenta ellipse), and shape the temporal pattern of sphingolipid accumulation. In the absence of Gba1b, sphingolipid levels are high across the day, promoting retraction and preventing growth, locking sLNv neurites in a reduced state. Perturbing *de novo* sphingolipid biosynthesis in sLNv using *Pdf>lace-RNAI* reduces sphingolipid accumulation, causing neurites to remain over-extended. Increasing Gba1b expression at night via *Pdf-GAL4* generates increased production of Cer sphingolipids in the morning, inverting the remodeling cycle. *p<0.05, ****p<0.0001, ANOVA, Tukey’s multiple comparisons. n > 25 sLNv neurites (A-I), n>30 flies (L-M), n>10 brains (N-O). Scale bar: 25μm (B-K). Data are represented as mean ± SEM. See also *Figure S7*, and *Table S1* for genotypes.

Given these observations and the circadian flux of sphingolipids in control brains (Figure 6D), we hypothesized that the cycle of growth and retraction of sLNv terminals might reflect cyclic changes in the relative balance of sphingolipid biosynthesis and catabolism. If such cyclic changes play an instructive role in determining when sLNv neurons grow and shrink, altering the timing of Gba1b expression could alter the growth cycle. To do this, we took advantage of the Pdf promoter, which increases transcriptional expression in a narrow interval at dusk (Figure S7N, S7P; Park et al., 2004). Strikingly, upon Gba1b induction in sLNv neurons with *Pdf-GAL4* in *gba1b^Δ^* mutants, cyclic remodeling was rescued but occurred antiphase to wild-type controls, with larger neurite spread at dusk and reduced spread at dawn (Fig 8F). As expected, lysosomal and aggregate phenotypes were not rescued in the rest of the brain (Figure S7J-K). These data show that Gba1b activity is sufficient to directly control the temporal phase of sLNv membrane remodeling.

To further test the hypothesis that sphingolipid flux is critical for sLNv remodeling, we impaired *de novo* sphingolipid biosynthesis using RNAi directed against Serine palmitoyltransferase (SPT/lace), the rate-limiting enzyme required for formation of sphingosine (which gives rise to Cer, GlcCer, CerPE, and higher-order sphingolipids (Figure 8G; Acharya and Acharya, 2005). Remarkably, sLNv neurons with reduced Lace activity were locked into a “morning-like” state, with an aberrantly large neurite volume that did not significantly cycle across the day (Figure 8H, Figure 8K). To test whether GlcCer is necessary for sLNv retraction, we knocked-down *Glucosylceramide Synthase (GlcT)*, the enzyme responsible for GlcCer synthesis, using *Pdf>GlcT^RNAi^*. Interestingly, sLNv neurons with reduced *GlcT* activity were modestly enlarged and also impaired in diurnal cycling (Figure 8 I-K), but less dramatically than the effects of lowering all sphingolipids via *lace* knockdown. Thus, cell-autonomous changes in sphingolipid biosynthesis and degradation impair structural plasticity in circadian circuits. Moreover, we infer that elevated levels of sphingolipids are associated with neurite shrinkage, while reduced levels of sphingolipids are associated with neurite growth. These data demonstrate that altering either sphingolipid biosynthesis or catabolism in sLNv circuitry is necessary and sufficient to alter sLNv neurite structure.

Consistent with sLNv cell state affecting behavior, both *Pdf>GlcT^RNAi^* and *Pdf>lace^RNAi^* flies had compromised activity profiles (Figure S7H, I), with *GlcT^RNAi^* increasing dawn activity and both *lace^RNAi^* and *GlcT^RNAi^* increasing evening activity. Intriguingly, there was a significant dampening of activity in *lace^RNAi^* flies at dusk when sLNv circuits are abnormally large. Remarkably, *Pdf-GAL4*>*Gba1b* expression in *gba1b^Δ^,* which inverts the cycle of sLNv remodeling, also triggered a similar dampening of activity at dusk when sLNv neurites are aberrantly large (Figure S7H-I). Thus, glia degradation and neuronal biosynthesis of specific sphingolipids are both necessary and sufficient to sculpt axon terminals across the day.

## Discussion

Our data demonstrate that glia produce Gba1b to non-autonomously control brain sphingolipids, protein degradation, and neurite remodeling in a circadian circuit (Figure 8N). We identified two specific glial subtypes, EG and PNG, as critical sources of Gba1b required for lysosomal function, proteostasis, circadian behaviors and neurite remodeling. While previous genetic studies in vertebrate models did not determine whether GBA was required in glia or neurons (Enquist et al., 2007)(Keatinge et al., 2015)(Uemura et al., 2015), lysosomal GBA expression in mice and humans is ~5-fold higher in glia and microglia compared to neurons (Zhang et al., 2016)(Zhang et al., 2014), and *gba* mice harbor aggregates in astrocytes (Osellame et al., 2013). Given these expression patterns and the striking similarities in brain phenotypes seen across *GBA* mutants in flies, fish, and mammals, there appears to be an evolutionarily ancient role for glia in regulating sphingolipid metabolism in the brain.

### Glia sculpt neurites to temporally control circuit remodeling

Previous work identified the cytoskeletal effector Rho1 and the transcription factor Mef2 as important for sLNv neurite remodeling (Petsakou et al., 2015)(Sivachenko et al., 2013). Our work demonstrates that these changes must be coordinated with membrane remodeling, particularly sphingolipid degradation and biosynthesis, both of which are required for membrane retraction and growth (Figure 8N). In control animals, sLNv neurites are enlarged at dawn, retract across the day to become stunted at dusk, and then re-extend during the night. This cycle coincides with a diurnal sphingolipid cycle in which specific GlcCer and Cer species are elevated during the retraction phase, and reduced during the growth phase (Figure 6E’, dusk-enriched species). In *gba1b^Δ^* mutants, the cycle of sLNv growth and retraction is blocked such that sLNv neurites remain in a stunted state across the day. In parallel, the sphingolipid cycle is also substantially blocked, and many sphingolipids are present at highly elevated levels. Conversely, in animals in which sphingolipid biosynthesis is globally reduced (*Pdf>lace^RNAi^)*, the cycle of sLNv remodeling is also blocked but sLNv neurites remain aberrantly extended throughout the day. Notably, Lace is bound by the transcription factor Clock (Abruzzi et al., 2011) and undergoes circadian changes in expression, increasing before dusk and peaking at midnight (Abruzzi et al., 2017)(Figure S7L), the same stage at which our RNAi perturbation would be expected to be strongest (given the Pdf promoter (Park et al., 2000)). Finally, changing the timing of Gba1b expression in sLNv neurites preserved the diurnal cycle of neurite growth and retraction, but inverted its phase such that sLNv neurites were reduced at dawn and enlarged at night. Taken together, these results demonstrate that carefully timed cycles of sphingolipid degradation and biosynthesis are instructive for the diurnal pattern of neurite remodeling.

Our lipidomics analysis demonstrates that the levels of Cer and GlcCer 14:1/18:1 and 14:1/20:1 species, as well as CerPE14:1/18:0 and CerPe 14:2/18:0, are elevated during the phase of sLNv neurite retraction, suggesting that these specific species might play an important regulatory role. Although sphingolipids are substantially less abundant than phospholipids (with Cer and GlcCer species representing <0.5% of neural membranes), their unique effects on membrane biophysics (Castro et al., 2014) (Carvalho et al., 2010) makes them well-poised to strongly affect membrane remodeling and structural plasticity. Finally, given that *GlcT* depletion (which blocks GlcCer but not Cer formation) did not increase sLNv volume compared to *lace* manipulations (which blocks GlcCer, Cer, and CerPE), we favor a model whereby Cer species drive membrane retraction. This hypothesis would be consistent with recent work revealing that diurnal changes in Cer species trigger retraction of microglia processes (Liu et al., 2021).

There is also strong evidence for sphingolipids regulating the cytoskeleton. For example, blocking *de novo* sphingolipid synthesis in fibroblasts acutely reduced cell area and lamellipodia formation (Meivar-Levy et al., 1997), and altered sphingolipid profiles were associated with shortened neurites and upregulated RhoA following *acid ceramidase* knockdown in neuroblastoma lines. Moreover, *GBA2* (non-lysosomal) knockout mice have cytoskeletal defects (Raju et al., 2015), shorter neurites, and dysregulate Rho GTPase localization (Woeste et al., 2019). Additionally, activity within larval LNv neurites triggers lipid-mediated structural alterations critical for circuit function dependent on Ceramide Synthase (Yin et al., 2018). Taken with our work, sphingolipids and their regulatory enzymes may function coordinately or even upstream to Rho1 and attendant cytoskeletal changes in sLNv remodeling, with more precise spatiotemporal control facilitated by glial degradation of remodeled neurite membranes.

While sLNv neurites are a dramatic example of membrane remodeling, many neurons grow and shrink across circadian cycles and varied environmental conditions (Heisenberg et al., 1995)(Barth et al., 1997)(Pyza and Meinertzhagen, 1999). Membrane turnover is likely important during structural plasticity at synapses, and indeed *gba1b* flies have memory defects (Davis et al., 2016). As EG and PNG are broadly distributed throughout the brain (Kremer et al., 2017), glia-mediated sphingolipid degradation may be central to membrane remodeling in many neural circuits. Interestingly, EG engulf neuronal membranes to clear damage in a sleep-dependent fashion (Doherty et al., 2009)(Musashe et al., 2016)(Stanhope et al., 2020), a cellular response potentially co-opted from a role in the daily remodeling of neurite membranes. Daily neurite remodeling is also a feature of the mouse suprachiasmatic nucleus (Becquet et al., 2008). Moreover, microglia, which express GBA in mice and humans, locally prune neurites during development and injury (Schafer et al., 2012)(Cangalaya et al., 2020), regulate sleep (Liu et al., 2021), and are enriched for sphingolipid catabolizing genes (Fitzner et al., 2020). Taken together, this suggests GBA could control many forms of structural plasticity across species.

### Protein aggregation is dynamically controlled by circadian state and neural activity

Our characterization of *gba1b* mutant animals revealed a surprising circadian cycle of protein aggregate accumulation and removal. We observed both activity-dependent and direct circadian control of aggregate burden. Protein aggregates have been described in wide-ranging disease models from flies to mice (Suresh et al., 2018) and are a prominent feature of neurodegeneration in humans (Ross and Poirier, 2004), including cells derived from *GBA* mutant mice and patients (Mazzulli et al., 2011)(Awad et al., 2015)(Schöndorf et al., 2014) (Osellame et al., 2013). Given our observations, it may prove critical to characterize aggregate accumulation (and lipid abundance) with respect to circadian time and neural activity. Indeed, circadian phagocytosis of amyloid-beta, a component of Alzheimer’s aggregates, was recently observed in cultured macrophages to be driven by circadian biosynthesis of heparan sulfate proteoglycans (Clark et al., 2022).

### Lipids as linchpins of disease: a central role for glia, circadian cycles, and sleep

Our data provide a direct mechanistic connection between the enzymatic activity of Gba1b in glia to circadian behavior through sLNv neurite remodeling, as well as broader effects on sleep behavior likely mediated by other circuits. Notably, *gba1b^Δ^* mutant animals display deficits in activity and sleep prior to overt accumulation of aberrant protein aggregates. Similarly, many Parkinson’s patients have sleep defects prior to development of clinical characteristics typically associated with neuropathological aggregate accumulation (Gan-Or et al., 2015)(Krohn et al., 2020). Many genes that impinge on lysosomal function and sphingolipid degradation are linked to Parkinson’s disease (Pan et al., 2008) (Lin et al., 2018), and glia are key regulators of Parkinson’s and other neurodegenerative diseases (Zuchero and Barres, 2015)(Liddelow et al., 2017). Moreover, precise control of lipid species is central to many neurodegenerative models and is often modulated by neural activity (Guttenplan et al., 2021)(Liu et al., 2015)(Tsai et al., 2019)(Jung et al., 2017) (Dasgupta et al., 2009) (Acharya et al., 2003) (Sellin et al., 2017). Recently, long-chain saturated lipids were found to mediate neurotoxicity by reactive astrocytes (Guttenplan et al, 2021), and we also observed increased nighttime long-chain saturated Cer/GlcCer species in *gba1b^Δ^* mutants (Table S4), which coincides with circadian aggregate burden. Based on these observations, mutations in lipid-regulating genes could impair glial remodeling of sleep circuits, and specific lipid species may diurnally drive cyclic aggregates. Sleep has been proposed to drive clearance of aggregates from the brain (Xie et al., 2013)(Zhang et al., 2018), and glia regulate sleep and circadian rhythms in flies and mice (Ng and Jackson, 2015) (Brancaccio et al., 2019)(Herrero et al., 2017). Moreover, cell-type specific functions for sphingolipids are known in glia (Ghosh et al., 2013)(Kunduri et al., 2018)(Dahlgaard et al., 2012), and regional lipid heterogeneity pervades the human brain (O’Brien and Sampson, 1965). Unraveling the complicated mechanisms of cell-type specific lipid synthesis and degradation may provide crucial insights into the connections between sleep, circadian rhythms, neurodegenerative diseases, and neuronal membrane dynamics.

### Limitations of the study

Our studies revealed that the brain is sensitive to the level of expression of Gba1b when doing rescue experiments, likely reflecting the fact that Gba1b itself is under tight transcriptional control and that very low levels of expression are sufficient to rescue most *gba1b* phenotypes (consistent with scRNA-seq data, Figure S3A). A key gap in the field is the absence of tools that afford both cell-type specificity and quantitative control of expression across physiologically relevant levels alongside precise temporal actuation. As such tools develop, further exploring the relative balance of sphingolipid biosynthesis and degradation in shaping neurite growth and retraction would be fascinating. In addition, while analyzing sLNv neurites at specific timepoints has provided key insights, being able to capture the dynamics of lipid turnover in a live imaging preparation would deepen this understanding. This requires the application of fluorescent probes for specific lipids that can be engaged cell-type specifically, combined with a chronic imaging preparation that can span both sleep and wake cycles (as has recently been described; Tainton-Heap et al., 2021)). Finally, while we have quantitatively measured changes in membrane lipid fluctuations across time in the entire brain using mass spectrometry, being able to measure changes in lipid pools from specific cell types, or even sub-cellular structures like neurites, would likely increase the strength and specificity of these signals, thereby providing new mechanistic insights into dynamic membrane remodeling.

## Acknowledgments

Research done at Stanford is conducted on Muwekma Ohlone land. We thank members of the Clandinin lab, Talbot lab, Zuchero lab, and Berfin Azizoğlu for comments on the manuscript; Minseung Choi and Yukun Alex Hao for analysis advice; Estela Stevenson for technical assistance; and Tom Braukmann for assistance with the DAM computer. We are grateful to Gabor Juhaz, Justin Blau, Michael Rosbash, Dragana Rogulja, Marc Freeman, Thomas Montine, Gerald Rubin, Nicholas Stavropoulos, Amita Sehgal, and Leo Pallanck for sharing reagents. Other stocks were obtained from the Bloomington Drosophila Stock Center (NIH P40OD018537). We also gratefully acknowledge the enthusiastic support of Tom Montine and the Matloff Fund, which made this work possible. This work was supported by NIH R01EY022638 (TRC), the NSF Graduate Resource Fellowship Program (GRFP) (JV), the Stanford Developmental Biology and Genetics Graduate Training Grant (JV), and by the Stanford Vision Core grant, NIH P30EY026877 (TRC).

## Author contributions

Conceptualization: JV, ET, RJR, MTC, TRC

Methodology: JV, ET, PK, AB, IA, IMRS, TPR, JDM, TH, EOP, VM, MTC, TRC

Investigation: JV, ET, IMRS, PK

Visualization: JV, ET, AB, TRC

Funding acquisition: JV, IA, TRC

Project administration: JV, VM, MTC, TRC

Supervision: VM, MTC, RJR, TRC

Writing – original draft: JV

Writing – review & editing: JV, TRC

## Declaration of interests

The authors declare no competing interests

---

**Figure S1.**
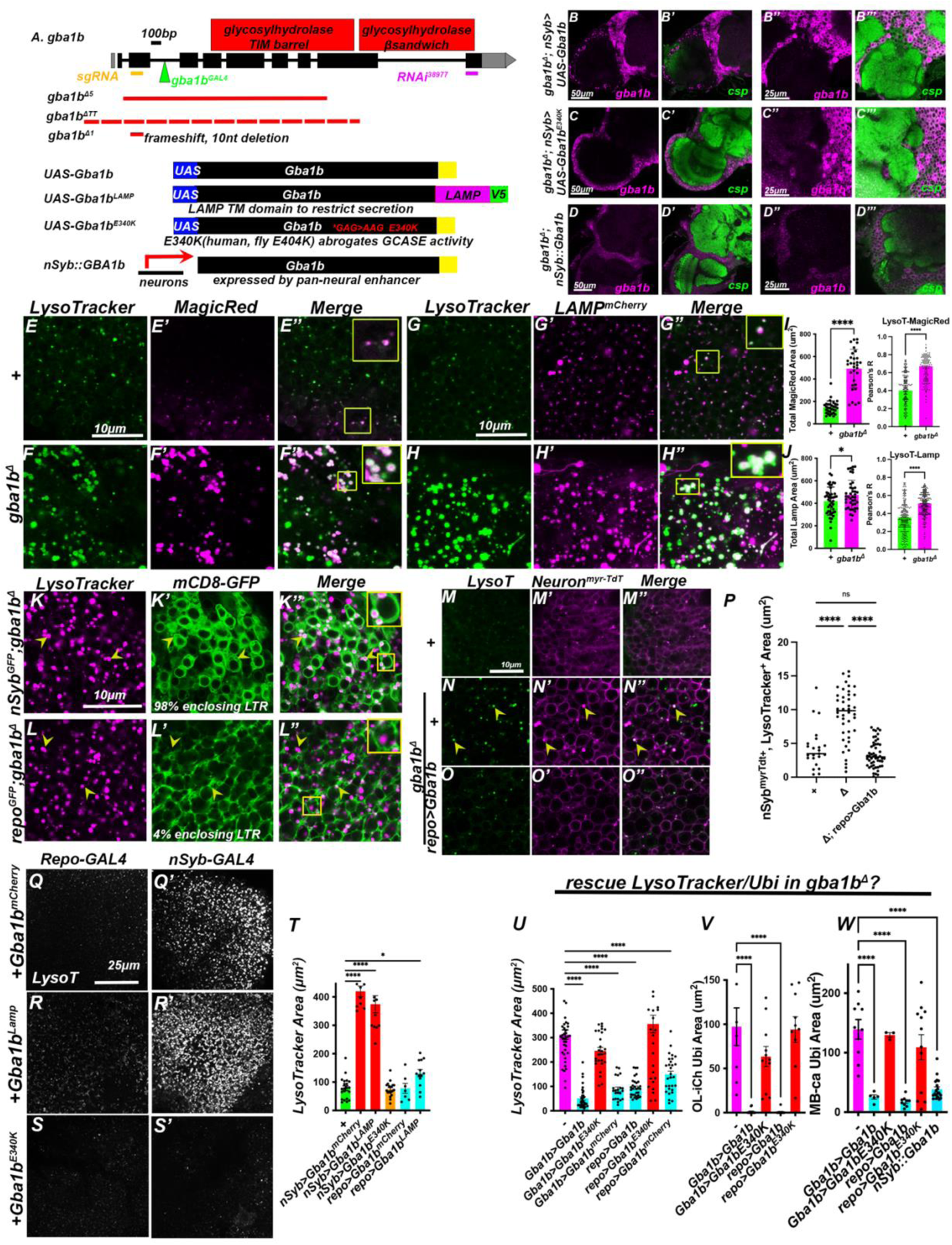
related to Figure 1. Gba1b tools, LysoTracker labels lysosomes in neurons, and rescue data. (A)The *Gba1b* locus (exons: black rectangles); target for cKO *sgRNA* (orange), *GBA^ΔGAL4^* (green; also terminates transcription and is a *Gba1b* null allele (Lee et al., 2018)), and *RNAi* (purple) are indicated. *gba1b* mutants used as transheterozygous combinations are outlined by dashed red lines. *Gba1b^ΔTT^* deletes the 5’ end of the coding sequence and *Qsox4* (not shown); *gba1b^Δ5^* flies also have a ~1.8kb insertion; *gba1b^Δ1^* generated a 10nt deletion and early frameshift (see supplemental table 1; deletes “CGATTATCCT”). We primarily used *gba1b^Δ1/TT^* heteroallelic combinations (to avoid analysis of flies carrying homozygous chromosomes) and designate this as *gba1b^Δ^*. Also schematized are the inducible *UAS-Gba1b* minigenes used for rescue, including *mCherry* C-terminal tagged, *LAMP*-transmembrane C-terminal tagged, and an *E340K* enzyme-dead version (following the nomenclature from the *GBA1* locus in humans; the fly equivalent site is E404). Along with *UA*S-overexpression constructs we made a *Gba1b* minigene directly under the control of a neural promoter (*nSyb::Gba1b*). (B-D) Anti-Gba1b (magenta) staining of *Gba1b* transgenes with neuropil counter-stained by Csp (green). (B) *UAS-Gba1b* and (C) *UAS-Gba1b^E340K^*expressed from neural promoters (*nSyb-GAL4*) in the *gba1b^Δ^*null background strongly induced anti-Gba1b signal in neural cortex in both optic lobe (B, B’, C, C’) and central brains (B’’, B’’’, C’’, C’’’). (D-D’’’) In contrast, *nSyb::Gba1b* expressed at much lower levels. (E-F) Optic lobe cortex co-stained for LysoTracker and Magic Red (marking Cathepsin B activity) in control (E) and *gba1b* mutants (F), with many degradative lysosomes present in *gba1b* mutants (inset). (G-H) Optic lobe cortex co-labeled for LysoTracker and *LAMP^mCherry^* in control (G) and *gba1b* mutant optic lobes (H). (I) Quantification of total area of Magic Red and colocalization analysis by Costes’ method (Costes et al., 2004) and Pearson’s-R. (J) Quantification of total area of *LAMP^mCherry^*and colocalization analysis by Costes’ method and Pearson’s-R. (K-L) mCD8*GFP* expressed in neurons or glia in *gba1b^Δ^*1-2 day old mutant brains revealed LysoTracker particles encircled by neural membrane but not glia membranes in the optic lobe cortex (arrowheads); 100% of neurons enclosed LysoTracker as opposed to 4% of glia, and 88% of Lysotracker could was inside neural-delimited CD8-GFP membranes. (M-P) Co-labeling distal optic lobe neural cortex membranes *(nSyb-LexA>LexAop-TdTom^myr^)* revealed small myr-Tdt puncta (magenta) that colocalized with Lysotracker (green). These neural-membrane+, LysoT+ puncta were larger in *gba1b* mutants (N), a size increase rescued by glial expression of Gba1b (*repo>Gba1b*) (O), quantified in P. (Q-T) Gain-of-function from neural *Gba1b*-overexpression. LysoTracker staining of optic lobe cortex in *UAS-Gba1b* overexpressing brains (in a wild-type background). Neural, but not glial, overexpression of *UAS-gba1b^mCherry^* (Q) or *UAS-Gba1b^LAMP^* (R) caused increased LysoTracker signal contingent on Gba1b enzyme activity as *UAS-Gba1b^E340K^* failed to cause a LysoTracker phenotype (S). (T) Quantification of LysoTracker data. (U-W) summary of *UAS-Gba1b* or *UAS-Gba1b^E340K^* rescue experiments in *gba1b^Δ^* by pan-glial (*repo-GAL4*) or *Gba1b-GAL4.* Shown are Lysotracker (U, optic lobe cortex from 2-5 day-old animals) and Ubi in Ol-ich (V) and MB-ca ((W), 15 day-old animals). Wild-type *UAS-Gba1b* but not *UAS-Gba1b^E340K^* can rescue *gba1b^Δ^* phenotypes when expressed by *repo-GAL4* or *Gba1b^GAL4^*. n>7 brains. Scale bar: 50μm (B-B’’, C-C’’, D-D’’), 25 μm (B’’-B’’’, C’’-C’’’, D’’-D’’’, Q-S’), and 10μm (E-M) *p<0.05, ****p<0.0001, ANOVA, Tukey’s multiple comparisons for normally distributed data, Kruskal-Wallis for nonparametric data (V-W). Data are represented as mean ± SEM. See *Table S1* for genotypes.

**Figure S2.**
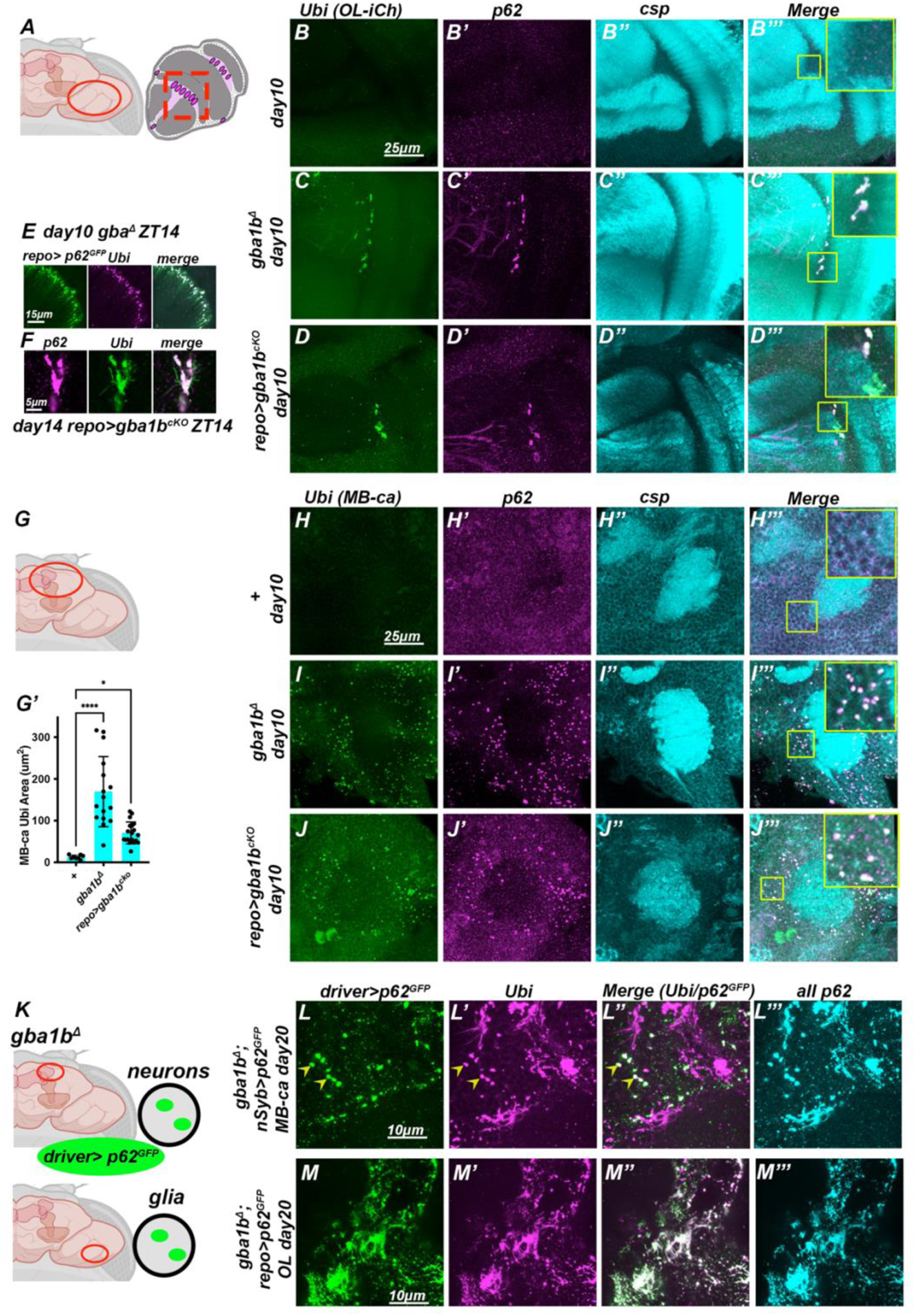
related to Figure 1. Diverse, stereotyped Ubi+, p62+ aggregates pervade gba1b^Δ^ brains. (A) Schematic of optic lobe inner chiasm (Ol-iCh). (B-D’’’) OL-iCh stained for Ubi (green), p62 (magenta), and neuropil (Csp, cyan). (H-J’’’) MB-ca stained for Ubi (green), p62 (magenta), and neuropil (Csp, blue). (B-B’’’) Control brain, day 10. (C-C’’’) *gba1b^Δ^* mutant brain, day 10. (D-D’’’) *repo>gba1b^cKO^* mutant brain, day 10. Insets show both *gba1b* perturbations cause chiasm aggregates. (E) Co-localization of OL-iCh Ubi aggregates with glial-expressed *P62-GFP* (Pircs et al., 2012)in *gba1b^Δ^*. (F) High-resolution image of a chiasm Ubi+p62+ aggregate from *repo>gba1b^cKO^*OL-iCh. (G) Schematic of anatomical location of MB-ca. (H-H’’’) Control brain MB-ca stained for aggregates at day 10. (I-I’’’) *gba1b^Δ^* mutant brain MB-ca stained for aggregates at day 10. (J-J’’’) *repo>gba1b^cKO^*mutant brain MB-ca stained for aggregates at day 10. Insets show both *gba1b* perturbations cause punctate aggregates in MB-ca cortex, quantified in G’. (K) schematic for detecting cell-type specificity of aggregates. (L-M) *UAS-p62^GFP^* expressed under the control of either neural or glial *GAL4* in 20-day old *gba1b^Δ^* mutant brains, stained with Ubi (magenta), GFP (green), and total p62 (cyan). (L-L’’’) Neural-expressed *p62-GFP* localized to puncta in MB-ca (arrowheads) but not more elaborate Ubi aggregate structures. (M-M’’’) glial-expressed *p62-GFP* colocalizes extensively with optic lobe Ubi aggregates. *p<0.05, ****p<0.0001, ANOVA, Tukey’s multiple comparisons. n>15 brains. Data are represented as mean ± SEM. Scale bar: 25μm (A-C’’’, F-H’’’), 15μm (D), 5μm (E), 10μm (I-J’’’). See *Table S1* for genotypes.

**Figure S3.**
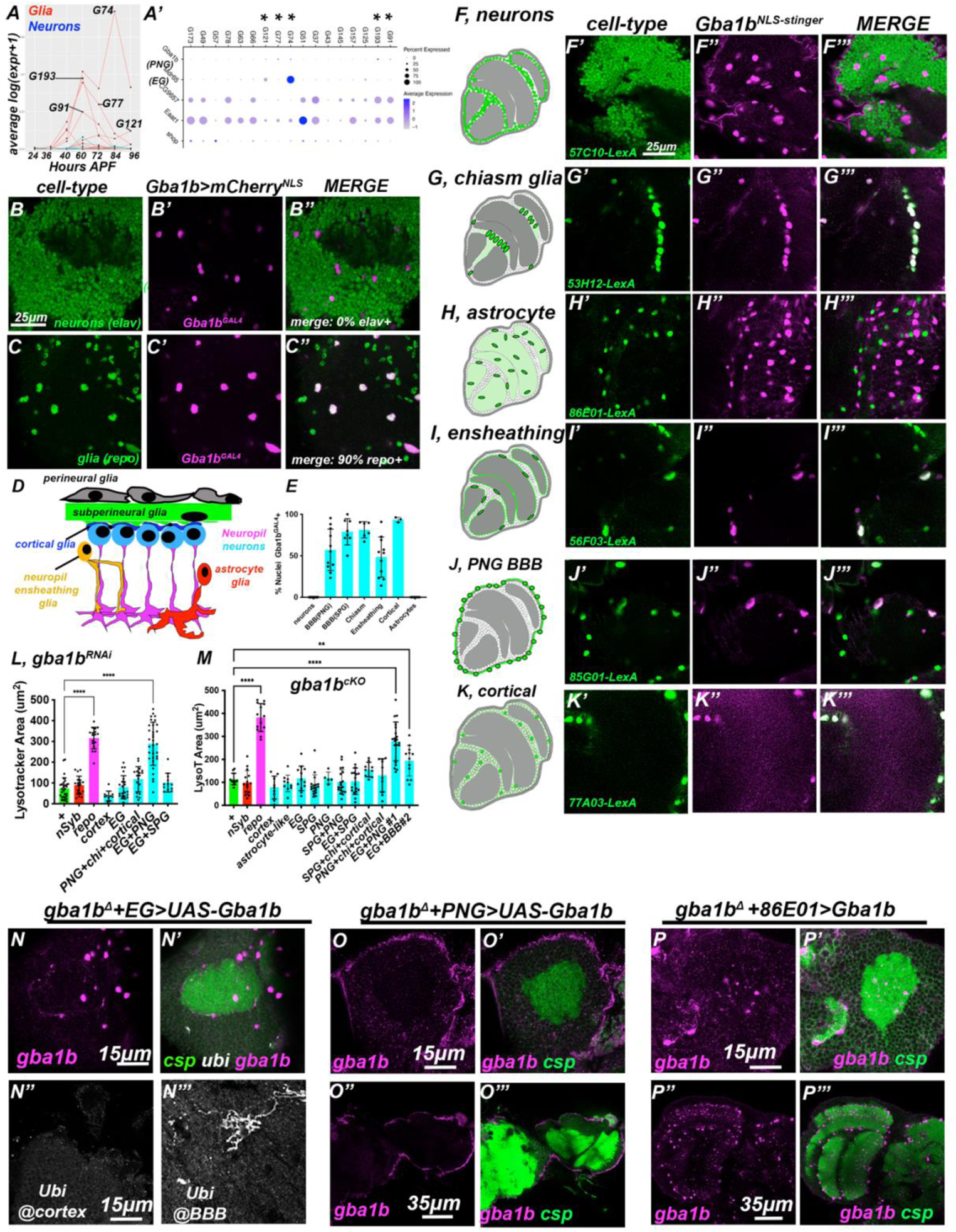
related to Figure 2. Gba1b is expressed in subsets of glia and required in both EG and PNG. (A) Developmental scRNA-seq data from (Kurmangaliyev et al, 2020) showing multiple clusters of glial-cells expressed Gba1b from 48 hours after pupal formation (APF) through 96 hours after pupal formation (red lines denote glia, blue lines denote neurons (the majority of which do not express Gba1b)). (A’’) While precise glial subtypes are not defined in this dataset, G74 expresses the PNG-enhancer *mdr65* and G91/G193 express the EG-enhancer CG*9657*. Asterisks indicate cell-types from (A); note that Gba1b expression (top row) is extremely low compared to these other genes (B) *gba1b^GAL4^* gene trap (Lee et al., 2018) drives expression of *UAS-mCherry^NLS^* (magenta), which never colocalized with anti-elav (neurons, green) but (C) extensively colocalized with anti-repo (glial, green). The repo negative, gba1b+ cells appeared to be tracheal cells (data not shown). (D) Schematic of different glia subtypes viewed in cross-section. (E) Quantification of colocalization of Gba1b expressing cells with different brain cells from (F-K). Cell-type specific orthogonal *LexA* nuclear marker (green) expression in the *Gba1b^GAL4^* (magenta) background. Schematics show cell body location and neuropil tiling in green. *GMR-LexA* driver designations are included in bottom left; all data are slices from optic lobes. (F-F’’’) Neurons. (G-G’’’) Chiasm glia. (H-H’’’) Astrocyte-like glia. (I-I’’’) Neuropil ensheathing glia. (J-J’’’) Perineural glia. (K-K’’’) Cortical glia. (L) Quantification of LysoTracker staining of optic lobe cortex from *gba1b^RNAi^* driven by both single *GAL4* drivers and various *GAL4* combinations. Only *EG* and *PNG* drivers together (*GMR56F03-GAL4; GMR85G01-GAL4)* displayed phenotypes comparable to pan-glial knockdown (*repo-GAL4*). (M) Quantification of LysoTracker from additional glia *GAL4* subtypes expressing somatic *CRISPR* (*gba1b^cKO^*), with an additional *EG+PNG* combination (*9137-GAL4; GMR83E12-GAL4*) that caused a LysoTracker phenotype. (N-N’) *EG-GAL4>Gba1b* expresses Gba1b (magenta) in cell bodies and around the MB-ca neuropil (Csp, green) and rescues cortex Ubi (white) but fails to rescue blood-brain barrier Ubi (N’’-N’’’). (O-O’’’) *PNG-GAL4>Gba1b* expresses Gba1b cell-autonomously in barrier cells in *gba1b^Δ^*, as indicated by anti-Gba1b (magenta) counterstained against neuropil (Csp, green) in CB (O-O’) and OL (O’’-O’’’). (P-P’’’) Astrocyte-expression of Gba1b induces strong Gba1b expression (magenta) in astrocyte cell bodies and processes in CB (P-P’) and OL (P’’-P’’’). *p<0.05, ****p<0.0001, ANOVA, Tukey’s multiple comparisons. Scale bar: 25μm (A-K) and 15μm (MB-ca) or 35μm (O-Q). n>5 optic lobes. See *Table S1* for genotypes.

**Figure S4.**
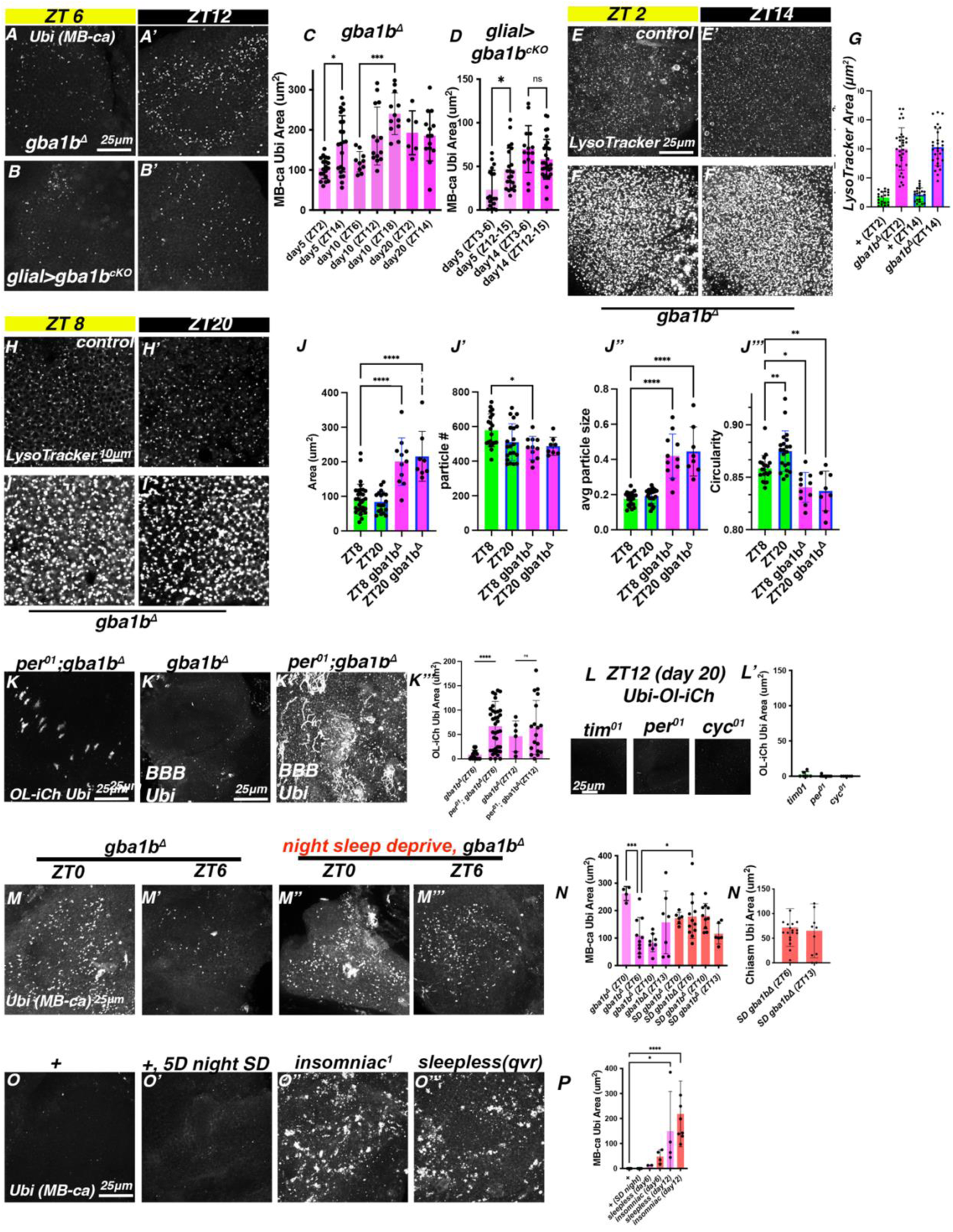
related to Figure 4. Additional circadian and sleep effects on lysosome size and Ubi aggregates. (A-A’) MB-ca Ubi structures in *gba1b^Δ^* or *repo>gba1b^cKO^* mutants (B-B’) have higher aggregate burden at ZT12 than ZT6. This effect is less apparent in older animals (darker magenta, C-D). (E-F) 5 day old control and *gba1b^Δ^*optic lobes analyzed for LysoTracker at daytime (ZT2 or ZT8) and nighttime (ZT14 or ZT20); max-intensity projections did not reveal substantial differences across time, with consistent elevation in *gba1b^Δ^* relative to controls (quantified in G). (H-I) High-resolution images of LysoTracker staining were used for particle analysis. (J) Particle analysis revealed subtle differences in control lysosomal populations across ZT: daytime control lysosomes had less circular morphology than nightime. Lysosome morphology in *gba1b^Δ^*mutants did not change diurnally, but were larger and less circular than controls. (K) 7 day-old, *per^01^; gba1b^Δ^* double mutants fail to clear Ubi at ZT6 and have higher Ubi burden in the BBB than *gba1b^Δ^* at ZT6. (L) OL-iCh of 20 day-old circadian clock mutants at ZT12 do not have prominent Ubi structures. (M) MB-Ca Ubi from *gba1b^Δ^* 6 day old controls were absent at ZT6 but accumulated following 2 nights of sleep-deprivation (SD, ZT12-ZT0), quantified in N. (O) Nighttime sleep deprivation does not trigger Ubi MB-Ca aggregates but short-sleeping mutants *qvr* and *insomniac* progressively accumulate Ubi aggregates (quantified in P). *p<0.05, ****p<0.0001, ANOVA, Tukey’s multiple comparisons for normally distributed data, Kruskal-Wallis test for nonparametric data (K). n>15 (A-L). n>5 (M-Q). Scale bar: 25μm except for H (10μm). Data are represented as mean ± SEM. See *Table S1* for genotypes.

**Figure S5.**
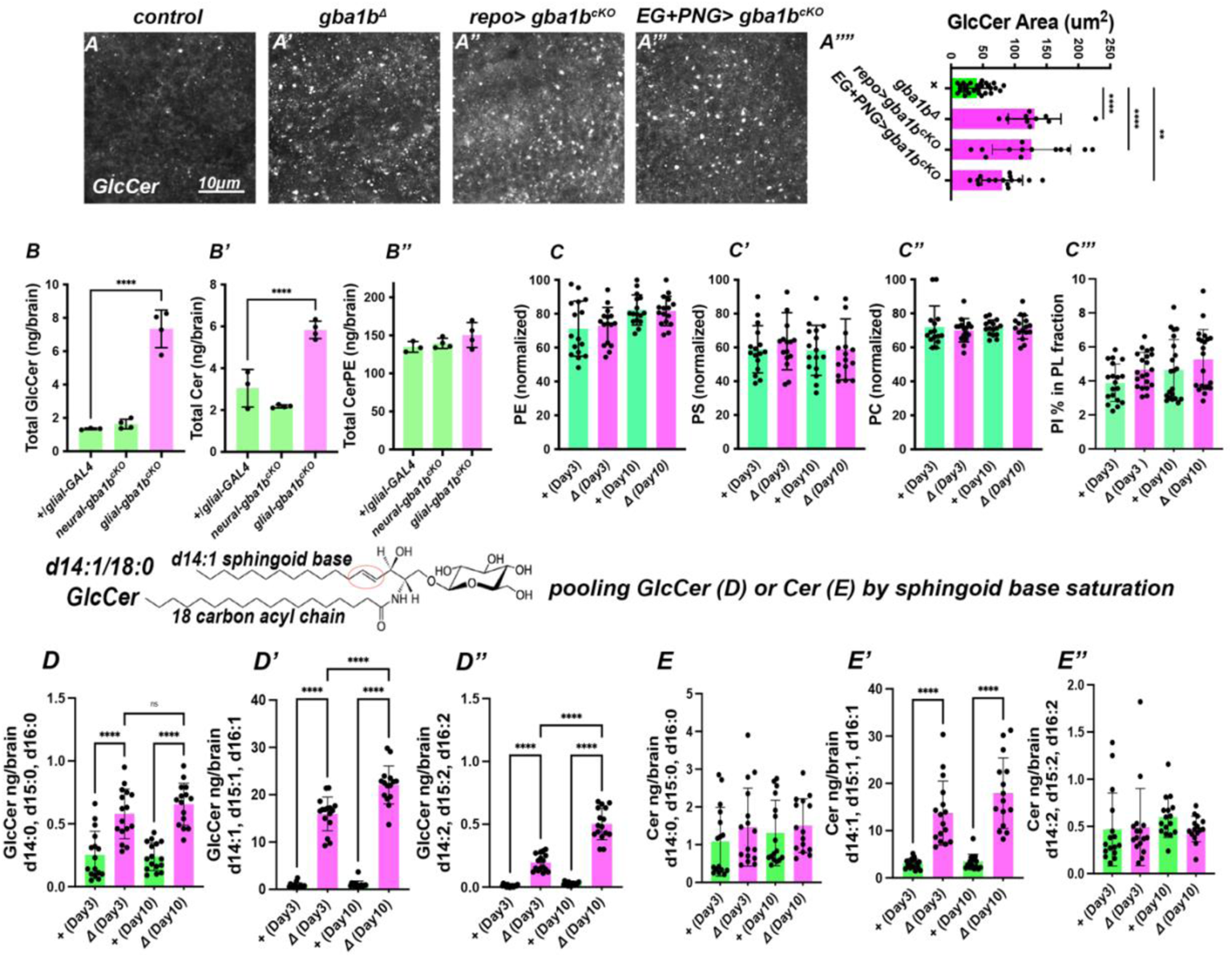
related to Figure 5. Further lipidomics analyses. (A) Staining against GlcCer in control (A), *gba1b^Δ^* (A’), *repo>gba1b^ckO^* (A’’), and *EG* and *PNG>gba1b^cKO^*at ZT12 in 10 day-old optic lobe cortex. Glial removal of *gba1b* triggered increased GlcCer puncta. (B) somatic *CRISPR* removal of gba1b (*gba1b^cKO^*) in glia (*repo*-*GAL4*) but not neurons (*nSyb-GAL4*) caused increased GlcCer and Cer in 10 day old brains. (C) Normalized phospholipid lipid abundance for controls (green) and *gba1b^Δ^* mutants (magenta) are plotted by age across two independent experiments. (C) Total normalized Phosphatidylethanolamine (PE). (C’) Total normalized Phosphatidylserine (PS). (C’’) Total normalized Phosphatidylcholine (PC). (C’’’) Total Phosphatidylinosital (PI). Bulk phospholipid levels were unaffected by *gba1b* removal. (D-E) GlcCer or Cer species analysis pooled by sphingoid base saturation (number of double bonds), independent of fatty acyl chain composition. (D, E) Saturated GlcCer and Cer species. (D’,E’) Monounsaturated GlcCer and Cer species. (D’’, E’’) Diunsaturated GlcCer and Cer species. While all GlcCer species were upregulated when assayed pooled by sphingoid base, both unsaturated (dihydroceramide, E) and diunsaturated Cer species (E’’) were unaffected by *gba1b^Δ^*. *p<0.05, ****p<0.0001, ANOVA, Tukey’s multiple comparisons for normally distributed data. n=15 brains/point, 8 biological replicates. Data are represented as mean ± SEM. See also *Table S1* for genotypes, and *Table S3* for raw lipidomics data.

**Figure S6.**
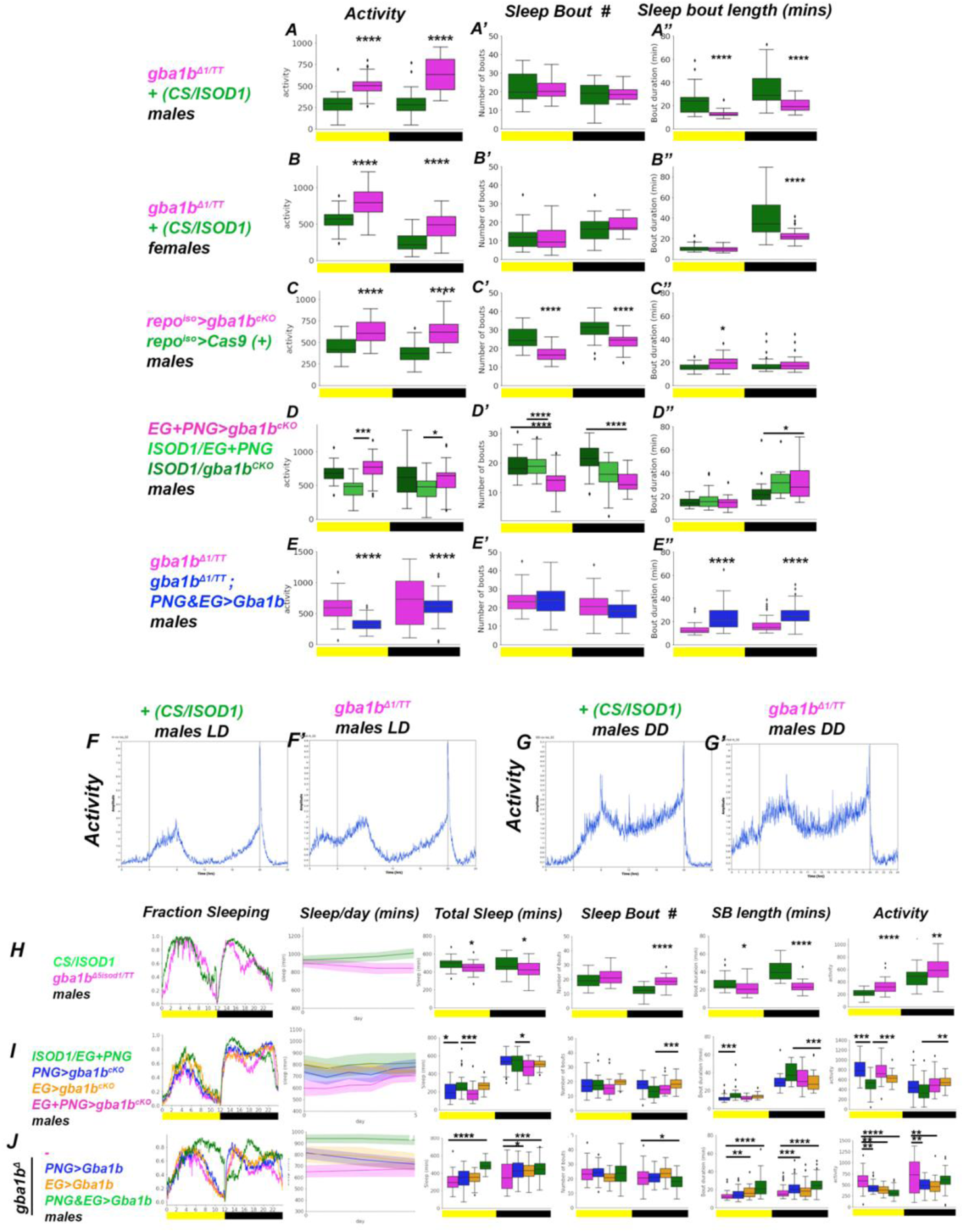
related to Figure 7. Additional Sleep Analysis. Activity, sleep bout number and sleep bout length (from day 5 to day 10 on LD) for the same genotypes in Figure 7. (A) *gba1b^Δ^* male flies (magenta) versus *CS/ISOD1* controls (green). (B) *gba1b^Δ^* female flies (magenta) versus *CS/ISOD1* controls. (C) Glial *gba1b^cKO^* males (magenta) versus control males (repo>cas9). (D) EG+PNG *gba1b^cKO^* males (magenta) versus control males (ISOD1/EG+PNG, and ISOD1/ *gba1b^cKO^*). (E) *gba1b^Δ^* males (magenta) versus rescue (EG+PNG>Gba1b) males (blue). (A, B, C, D, E) Activity. (A’, B’, C’, D’, E’) Sleep bout number. (A’’, B’’, C’’, D’’, E’’) Sleep bout length. *gba1b^Δ^, repo>gba1b^cKO^* and *EG+PNG>gba1b^cKO^* have higher total activity, and *repo>gba1b^cKO^* and *EG+PNG>gba1b^cKO^* have fewer sleep bouts. (F-G) Actograms of *CS/ISO* and *gba1b^Δ^* males on LD cycles (F) and dark-dark (DD) cycles (G-G’). Peaks of morning and evening activity occur in both cases, though the strength of *gba1b* rhythmicity was reduced, see Table S5. (H) *gba1b^Δ5/TT^*males (magenta) sleep less than control males (green). (I) Removing Gba1b in solely EG (orange) or PNG (blue) using somatic *CRISPR* (*gba1b^cKO^*) did not cause as strong a sleep defect as dual removal in both EG+PNG (magenta, data from Figure 7). (J) Rescuing Gba1b in *gba1b^Δ1^*(magenta) in single glia types (EG (orange) or PNG (blue)) did not restore sleep behavior as fully as EG+PNG dual-rescue of Gba1b (green, dual rescue from Figure 7). *p<0.05, ****p<0.0001, ANOVA, Tukey’s multiple comparisons or Welch’s ANOVA depending on equality of variance. For nonparametric sleep bouts, Mann-Whitney U test was used. n>28 flies. Data are represented as mean ± SEM for activity and bout boxplots, each showing the IQR. See *Table S1* for genotypes.

**Figure S7.**
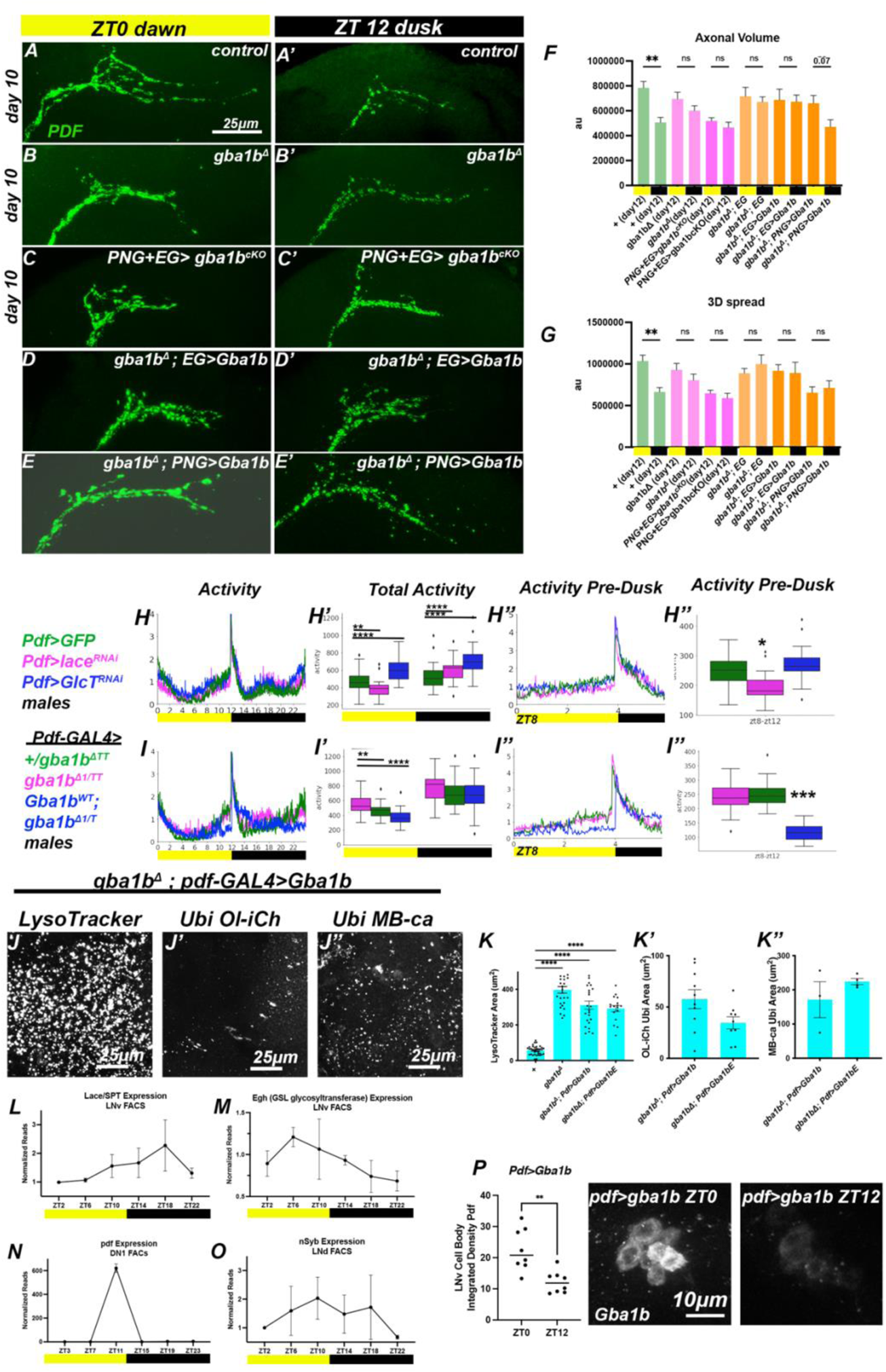
Related to Figure 8. sLNv remodeling and *Pdf-GAL4* rescue data. Quantification of sLNv morphology by anti-PDF in 10 day-old flies. (A) Control. (B) *gba1b^Δ^*. (C) *PNG+EG>gba1b^cKO^* (D). (D) Rescue of *gba1b^Δ^* using EG-Gal4. (E) Rescue of *gba1b^Δ^* using PNG-Gal4. (F-G) Quantification of 3D-spread and axonal volume. Rescue experiments using either the EG Gal or PNG Gal4 alone did not rescue *gba1b^Δ^*. (H-I) DAM data from day 5 to day 10 adult males. (H) *Pdf>GlcT^RNAi^*(blue), *Pdf>lace^RNAi^* (magenta), and *Pdf>GFP* control (green) activity profiles, quantified in (H’). (H’’-H’’’). Focusing on activity pre-dusk (ZT8-12), when sLNv neurites are shrinking. (I) *pdf-GAL4* > *Gba1b* activity profiles (blue) in *gba1b^Δ^* mutants are shifted from both controls (green) and *gba1b^Δ^*mutants (magenta), quantified in (I) across the entire day and at ZT8-12 (I’’-I’’)). (J-K) *gba1b^Δ^*mutants expressing Gba1b via *Pdf-GAL4* do not rescue LysoTracker or Ubiquitin in the Ol-iCh or MB-ca (quantified in J-K). (L-O) Data replotted from (Abruzzi et al, 2017), where various clock neurons (LNv, DN1, LNd) were FACS-sorted and analyzed across ZT by RNA-sequencing. This dataset identified both *lace/SPT*, the rate-limiting step in sphingolipid biosynthesis, and *egghead (egh)*, a glycosyltransferase critical for making higher-order glycosphingolipids, as cycling transcripts within LNv cells. We plotted the average of this data normalized to ZT2 across two independent experiments, which revealed a strong midnight peak of *lace* expression (L). (M) *egh* expression increases following *lace* expression and is higher in the morning and midday. (N) *pdf* has an large peak of expression at ZT11, and PDF protein levels show corresponding increase through night to dawn (Park et al., 2000). (O) *nSyb* was identified as weakly cycling by RNA-seq and is Clock-bound (Abruzzi et al., 2011). (P) *Pdf>Gba1b* sLNv cells stained with anti-Gba1b shows strong circadian fluctuations in protein *p<0.05, ****p<0.0001, ANOVA, Tukey’s multiple comparisons or Welch’s ANOVA depending on equality of data variance. For nonparametric sleep bouts Mann-Whitney U test was used. n >24 sLNv neurites (A-E), n>30 flies (H-I), n>3 (K), n>5 (P). Scale bar: 25μm (A-E, J-K), 10μm (P). Data are represented as mean ± SEM See *Table S1* for genotypes.

## STAR Methods

### KEY RESOURCES TABLE

| REAGENT or RESOURCE | SOURCE | IDENTIFIER |
| --- | --- | --- |
| Antibodies |  |  |
| Gba1b (anti-rabbit polyclonal) | This paper, abclonal | Anti-Gba1b |
| Csp (anti-mouse IgG2b) | Developmental Studies Hybridoma Bank | Cat#DCSP-2 (6D6)<br>RRID: AB_528183 |
| Ubi (anti-mouse IgG1) | Enzo Life Sciences | Cat# BML-PW8810,<br>RRID:AB_10541840 |
| p62 (anti-rabbit polyclonal) | Abcam | Cat# ab178440 |
| GFP (anti-rabbit polyclonal) | Abcam | Cat# ab13970 |
| GlcCer (anti-rabbit polyclonal) | Glycobiotech GmbH | Cat# RAS_0010 |
| PDF (anti-mouse IgG2b) | Developmental Studies Hybridoma Bank | Cat# PDF C7,<br>RRID: AB_760350) |
| Bruchpilot (anti-mouse IgG1) | Developmental Studies Hybridoma Bank | Cat# nc82;<br>RRID: AB_2314866 |
| Goat anti-Chicken IgY (H+L) Secondary Antibody, Alexa Fluor 488 | moFisher Scientific | Cat# A-11039,<br>RRID:AB_2534096) |
| Goat anti-Rabbit IgG (H+L) Cross-Adsorbed Secondary Antibody, Cyanine3 | ThermoFisher Scientific | Cat# A10520,<br>RRID:AB_2534029 |
| Goat Anti-Mouse IgG2b Antibody, Alexa Fluor 488 Conjugated | ThermoFisher Scientific | Cat# A-21141,<br>RRID:AB_141626) |
| Goat anti-Mouse IgG1 Cross-Adsorbed Secondary Antibody, Alexa Fluor 647 | ThermoFisher Scientific | Cat# A-21240, RRID:AB_2535809) |
| Goat anti-Mouse IgG2b Secondary Antibody, Alexa Fluor 633 | ThermoFisher Scientific | Cat# A-21146, RRID:AB_2535782) |
| Critical commercial assays |  |  |
| LysoTracker Red DND-99 | ThermoFisher Scientific | Cat# L7528 |
| LysoTracker Deep Red | ThermoFisher Scientific | Cat# L12492 |
| LysoTracker™ Green DND-26 | ThermoFisher Scientific | Cat# L7526 |
| MagicRed Cathepsin Assay | Abcam | Cat# ab270772 |
| Nile Red | ThermoFisher Scientific | Cat# N1142 |
| Experimental models: Organisms/strains |  |  |
| *D. melanogaster*: +(ISOD1) | Clandinin lab | N/A |
| *D. melanogaster*: +(CantonS) | Clandinin lab | N/A |
| *D. melanogaster*: $P\{y[+t7.7]v[+t1.8]=TRiP.HMS01893\}attP40$ | Bloomington Drosophila Stock Center | BDSC#38977, FBst0038977 |
| *D. melanogaster*: $gba1b[\Delta TT]$ | Davis et al., 2016 | N/A |
| *D. melanogaster*: $gba1b[\Delta 1]$ | this paper | N/A |
| *D. melanogaster*: $gba1b[\Delta 5]$ | Perlara (Joshua Mast) | N/A |
| *D. melanogaster*: $P\{y[+t7.7]v[+t1.8]=TKO.GS00895\}attP40$ | Bloomington Drosophila Stock Center | BDSC#77100, FBst0077100 |
| *D. melanogaster*: $P\{y[+t7.7]w[+mC]=UAS-Cas9.P2\}attP2/TM6B, Tb[1]$ | Bloomington Drosophila Stock Center | BDSC#58986, FBst0058986 |
| *D. melanogaster*: $nSyb::Gba1b\{attP16\}$ | this paper | N/A |
| *D. melanogaster*: $UAS-gba1b-LAMP-V5\{attP16\}$ | this paper | N/A |
| *D. melanogaster*: $UAS-gba1b\{attP16\}$ | this paper | N/A |
| *D. melanogaster*: $UAS-gba1b\{attP2\}$ | this paper | N/A |
| *D. melanogaster*: $UAS-gba1b[E340K]\{attP2\}$ | this paper | N/A |
| *D. melanogaster*: $UAS-gba1b[E340K]\{attP16\}$ | this paper | N/A |
| *D. melanogaster*: $UAS-gba1b-mCherry\{attP16\}$ | this paper | N/A |
| *D. melanogaster*: $UAS-gba1b-mCherry\{attP2\}$ | this paper | N/A |
| *D. melanogaster*:<br>*TI{GFP[3xP3.cLa]=CRIMIC.TG4.0}gba1b1b[CR00541-TG4.0]/TM3, Sb[1] Ser[1]* | Bloomington<br>Drosophila Stock<br>Center | BDSC#78943,<br>FBst0078943 |
| *D. melanogaster*: *P{y[+t7.7]*<br>*w[+mC]=8XLexAop2-FLPL}attP40* | Bloomington<br>Drosophila Stock<br>Center | BDSC#55820,<br>FBst0055820 |
| *D. melanogaster*: *P{13XlexAop-myr::tdTomato}attP2* | Gerald Rubin | FBtp0093486 |
| *D. melanogaster*: *w[\*]; P{w[+mC]=UAS-Stinger}2, PBac{y[+mDint2]*<br>*w[+mC]=13XLexAop2-IVS-tdTomato.nls}VK00022* | Bloomington<br>Drosophila Stock<br>Center | BDSC#66680,<br>FBst0066680 |
| *D. melanogaster*: *w[\*]; P{w[+mC]=UAS-mCherry.NLS}2; MKRS/TM6B, Tb[1]* | Bloomington<br>Drosophila Stock<br>Center | BDSC#38425,<br>FBst0038425 |
| *D. melanogaster*: *P{10XUAS-IVS-mCD8::GFP}attP2* | Bloomington<br>Drosophila Stock<br>Center | BDSC#32185,<br>FBst0032185 |
| *D. melanogaster*: *P{y[+t7.7]*<br>*w[+mC]=GMR57C10-GAL4}attP2* | Bloomington<br>Drosophila Stock<br>Center | BDSC#39171,<br>FBst0039171 |
| *D. melanogaster*: *P{y[+t7.7]*<br>*w[+mC]=GMR57C10-lexA}attP40/CyO* | Bloomington<br>Drosophila Stock<br>Center | BDSC#52817,<br>FBst0052817 |
| *D. melanogaster*: *w[1118]; P{y[+t7.7]*<br>*w[+mC]=GMR77A03-GAL4}attP2* | Bloomington<br>Drosophila Stock<br>Center | BDSC#39944,<br>FBst0039944 |
| *D. melanogaster*: *w[1118]; P{y[+t7.7]*<br>*w[+mC]=GMR77A03-lexA}attP40* | Bloomington<br>Drosophila Stock<br>Center | BDSC#54108,<br>FBst0054108 |
| *D. melanogaster*: *w[1118]; P{y[+t7.7]*<br>*w[+mC]=GMR85G01-lexA}attP40* | Bloomington<br>Drosophila Stock<br>Center | BDSC#54285,<br>FBst0054285 |
| *D. melanogaster*: *w[1118]; P{y[+t7.7]*<br>*w[+mC]=GMR85G01-GAL4}attP2* | Bloomington<br>Drosophila Stock<br>Center | BDSC#40436,<br>FBst0040436 |
| *D. melanogaster*: *w[1118]; P{y[+t7.7]*<br>*w[+mC]=GMR54C07-GAL4}attP2* | Bloomington<br>Drosophila Stock<br>Center | BDSC#50472,<br>FBst0050472 |
| *D. melanogaster*: *w[1118]; P{y[+t7.7]*<br>*w[+mC]=GMR54C07-lexA}attP40* | Bloomington<br>Drosophila Stock<br>Center | BDSC#61562,<br>FBst0061562 |
| *D. melanogaster*: *w[1118]; P{y[+t7.7]*<br>*w[+mC]=GMR53H12-GAL4}attP2* | Bloomington<br>Drosophila Stock<br>Center | BDSC#50456,<br>FBst0050456 |
| *D. melanogaster*: *w*[1118]; *P*{*y*[+*t*7.7]<br>*w*[+*mC*]=*GMR53H12-lexA*}*attP40* | Bloomington<br>Drosophila Stock<br>Center | BDSC#53573,<br>FBst0053573 |
| *D. melanogaster*: *w*[1118]; *P*{*y*[+*t*7.7]<br>*w*[+*mC*]=*GMR56F03-GAL4*}*attP2* | Bloomington<br>Drosophila Stock<br>Center | BDSC#39157,<br>FBst0039157 |
| *D. melanogaster*: *w*[1118]; *P*{*y*[+*t*7.7]<br>*w*[+*mC*]=*GMR56F03-lexA*}*attP40* | Bloomington<br>Drosophila Stock<br>Center | BDSC#53574,<br>FBst0053574 |
| *D. melanogaster*: *w*[*]; *P*{*y*[+*t*7.7]<br>*w*[+*mC*]=*GMR56F03-GAL4*}*attP24/CyO* | Bloomington<br>Drosophila Stock<br>Center | BDSC#77469,<br>FBst0077469 |
| *D. melanogaster*: *w*[*]; *P*{*w*[+*mC*]=*nrv2-*<br>*GAL4.S*}3; *P*{*nrv2-GAL4.S*}8 | Bloomington<br>Drosophila Stock<br>Center | BDSC#77469,<br>FBst0077469 |
| *D. melanogaster*: *w*[1118]; *P*{*y*[+*t*7.7]<br>*w*[+*mC*]=*GMR86E01-GAL4*}*attP2* | Bloomington<br>Drosophila Stock<br>Center | BDSC#45914,<br>FBst0045914 |
| *D. melanogaster*: *w*[1118]; *P*{*y*[+*t*7.7]<br>*w*[+*mC*]=*GMR86E01-lexA*}*attP40/CyO* | Bloomington<br>Drosophila Stock<br>Center | BDSC#54287,<br>FBst0054287 |
| *D. melanogaster*: *w* [*]; *P* { *w* [+ *m* *]= *alrm-*<br>*GAL4.D* }3/ *CyO*; *Dr* [1]/ *TM3, Sb* [1] | Bloomington<br>Drosophila Stock<br>Center | BDSC#67031,<br>FBst0067031 |
| *D. melanogaster*: *w*[1118]; *P*{*y*[+*t*7.7]<br>*w*[+*mC*]=*GMR83E12-GAL4*}*attP2* | Bloomington<br>Drosophila Stock<br>Center | BDSC#40363,<br>FBst0040363 |
| *D. melanogaster*: *w*[1118]; *P*{*y*[+*t*7.7]<br>*w*[+*mC*]=*GMR83E12-lexA*}*attP40* | Bloomington<br>Drosophila Stock<br>Center | BDSC#54288,<br>FBst0054288 |
| *D. melanogaster*: *w* [*]; *P* { *w* [+ *m* *]= *moody-*<br>*GAL4.SPG* }2 | Bloomington<br>Drosophila Stock<br>Center | BDSC#90883,<br>FBst0090883 |
| *D. melanogaster*: *P* { *w* [+ *m* *]= *GAL4* } *repo/TM3,*<br>*Sb* [1] | Bloomington<br>Drosophila Stock<br>Center | BDSC#7415,<br>FBst007415 |
| *D. melanogaster*: *per*<sup>01</sup> | Bloomington<br>Drosophila Stock<br>Center | BDSC#80928,<br>FBst0080928 |
| *D. melanogaster*: +; *Pdf-GAL4*; + | Dragana Rogulja | FBtp0011844 |
| *D. melanogaster*: *LAMP*<sup>3xmCherry</sup> | Gábor Juhász | FBal0325101 |
| *D. melanogaster*: *inc*<sup>1</sup> | Nicholas<br>Stavropoulos | FBal0266013 |
| *D. melanogaster*: *inc*<sup>2</sup> | Nicholas<br>Stavropoulos | FBal0162225 |
| *D. melanogaster*: *qvr*<sup>EY04063</sup> (*sleepless*) | Bloomington Drosophila Stock Center | FBst0016588 |
| *D. melanogaster*: *P{GawB}9-1379-137* | Amita Sehgal | FBal0344513 |
| *D. melanogaster*: *P{GMR83E12-GAL4}* | Bloomington Drosophila Stock Center | FBtp0063377 |
| *D. melanogaster*: *y1 v1*; *P{TRiP.HMC03219}attP40 [lace-RNAi]* | Bloomington Drosophila Stock Center | FBti0157638 |
| *D. melanogaster* : *P{KK107304}VIE-260B [GlcT-RNAi]* | Vienna Drosophila Resource Center | FBst0479877 |
| Oligonucleotides |  |  |
| Primers for *Gba1b* locus sequencing, see Table S2 | This paper |  |
| Recombinant DNA |  |  |
| Sequences of *Gba1b* plasmids, see Table S2 | This paper |  |
| Software and algorithms |  |  |
| ImageJ (FIJI) | ImageJ | RRID: SCR_003070 |
| GraphPad Prism 9 | GraphPad | RRID: SCR_002798 |
| MATLAB_R2021a | MATLAB | RRID: SCR_001622 |
| Python 3.7.1 | Python | RRID: SCR_008394 |
| R Studio 1.3.1093 | R | RRID: SCR_000432 |
| Clocklab 2.1 | Actimetrics | RRID:SCR_014309 |

***Table S1, detailed genotypes and n***

***Table S2, sequences for primers and UAS-Gba1b plasmids***

***Table S3, lipidomics data sphingolipids and phospholipids (ng/brain)***

***Table S4, differentially expressed lipid subspecies based on time or genotype***

***Table S5, circadian rhythm strength and comparison of sleep software analyses***

***Table S6, circadian-regulated lipid enzymes from (Abruzzi et al, 2011; Abruzzi et al, 2017)***

### Materials and Methods

#### Lead contact

Further information and requests for resources and reagents should be directed to and will be fulfilled by the lead contact, Tom Clandinin.

#### Materials availability

Flies and plasmids are available from the lead contact, and flies will be deposited at Bloomington Drosophila Stock Center (BDSC).

#### Data and code availability

Sphingolipid lipidomics are deposited as Supplemental Table 3. Python code used for sleep analysis is available at https://github.com/ClandininLab/sleep_analysis. All FIJI and R scripts are available upon request. All data and information required to analyze the data in this paper are available from the lead contact upon request.

#### *Drosophila* Maintenance

Flies were maintained on standard molasses food. Freshly prepared food was used for all behavior, lipidomics, and circadian IHC/live imaging experiments. Stocks were maintained in light/dark (LD) incubators for 12:12 light:dark cycles at 24-25°C, for all experiments except those using *gba1b^RNAi^*, which were performed at 29°C. For dark-dark rescue experiments (DD), crosses were set in DD incubators wrapped in foil, and experimental F1s were only temporarily exposed to light every 3 days to change food. Females were used for all experiments, except as noted; however, no obvious sex differences were observed (data not shown). For circadian experiments with age-matched genotypes, F1 experimental genotypes were collected *en masse* and then split into independent vials housed longitudinally in the same incubator, on the same shelf (to ensure similar lighting). Individual vials were removed immediately prior to dissection.

#### Drosophila Genetics

See Table S1 for a complete list of genotypes and sample sizes. Unless otherwise stated, the control genotype was an isogenized lab-stock (*ISOD1, 75*) placed *in trans* to an independent control, *CantonS* (*CS*) (“*+*”, *ISOD1/CS*). The *gba1b^Δ1^* frameshift allele (Fig. S1A) was induced by crossing *nos-Cas9* (germline *Cas9*) to a ubiquitously expressed gba1b guide (BDSC#77100) targeting an early exon to generate males carrying *nos-Cas9/gba1b^sgRNA^*on the 2nd chromosome. These males carrying potential *gba1b^Δ?^* alleles were crossed to an isogenic *+;+;pr/TM6b* virgin stock, and 10 males (*+;+;gba1b^Δ??^/Tm6b*) were outcrossed to a double-balancer stock. F3 homozygotes were screened for LysoTracker phenotypes; 9/10 lines had strong *gba1b* loss-of-function LysoTracker enlargement and subsequent sequencing revealed deletions at the predicted guide locus of 1,3,6, and 10nt. The 10nt deletion, *gba1b^Δ1^*, was used *in trans* to *gba1b^ΔTT^*(Davis et al., 2016) (“*gba1b^Δ^*”, *gba1b^Δ^*). In certain experiments, *gba1b^Δ1^* was used in *trans* to an alternative *gba1b* allele, *gba1b^Δ5^*, which deletes most exons (but not the start codon). *gba1b^Δ5^* was isogenized by backcrossing 5 generations into the *ISOD1* background and recombined with *GAL4* lines on chromosome III. *repo-GAL4* was also isogenized by backcrossing 5 generations into *ISOD1* for *repo*>*gba1b^cKO^*sleep experiments. For all rescue assays with *gba1b* alleles, the presence of two *gba1b* alleles was confirmed by taking F1 siblings that failed to carry a component of the rescuing constructs (*GAL4/UAS* or otherwise) and checking for brain LysoTracker enlargement. Rescuing transgenes were screened for GAL4-independent rescue by checking *UAS-Gba1b; gba1b^Δ^* controls for *gba1b* loss-of-function phenotypes using LysoTracker and anti-Ubi staining.

#### Molecular Genetics

All *UAS-Gba1b* overexpression constructs were synthesized and cloned (Genscript) and injected into *attP* landing sites (BestGene; Table S2). *attp2* was used for all 3^rd^ chromosome inserts, while *attP16* and *attP30* were used for 2^nd^ chromosome, selected based on inducibilty and lower background expression than *attP40* (Markstein et al., 2008). The *pJRF7-20XUAS-IVS-mCD8::GFP* overexpression backbone harboring *attB* (Addgene # (26220, Pfeiffer et al., 2010)) was digested with XbaI and XhoI to remove intervening *mCD8-GFP*, which was replaced by a *gba1b* minigene (exons1-9 including signal sequence, omitting introns, stop codon, and UTRs); a Kozak sequence CAAA proceeds the ATG start codon before the *gba1b* minigene, and all constructs terminate with *3XSTOP* and a *SV40* terminator. In *UAS-Gba1b^mCherry^* constructs, a NotI site and *10X-Proline* linker (GCGGCCGCGAGGTGGCGGAGGTGGCGGAGGTGGCGGAGG) were inserted prior to the *mCherry* sequence harboring an 11 N-terminal nucleotide deletion to improve *mCherry* stability and prevent cleavage from Gba1b (Huang et al, 2014). For *UAS-Gba1b^V5^* constructs, a NotI site and 6XGly linker were inserted (GCGGCCGCGCAGCAGCAG) before a *V5* epitope tag. A cloning error caused a frameshift immediately after the NotI linker and rendered the V5 epitope unusable. For the *UAS-Gba1b^E340K^* enzyme-dead variant, the human-equivalent of *E340K*, *E404K* in *Drosophila* (AACACGAAGTCCTGC→ AACACGGAGTCCTGC), was induced by site-directed mutagenesis and validated by sequencing (Sequetech; see Table S2 for sequencing primers). For *UAS-Gba1b^LAMP^*, the *LAMP1* transmembrane sequence from *Drosophila* (GAAGCTCATGTAACCGCGGGAGGTGGAACGACGCCTAGCACACAGATATGATATTAATACCACCAAAATTAATGCAGCTAAAGCAATTCCAACGGCAATGGGAACCAC) was appended to the C-terminal of a *UAS-Gba1b* backbone, following a NotI-(GS)_5_-(V5)_3_ linker and epitope tag. For *nSyb::Gba1b,* the 824bp promoter used in the pan-neural *nSyb* enhancer *GMR57C10* was synthesized and cloned into *UAS-Gba1b^E340K^* by digesting with HindIII and BGIII, which deletes the *20XUAS* upstream (*5XGAL4-DBD*)_4_ sequences as well as the HS promoter but leaves the *mhc IVS* intact. This places the Gba1b minigene ~100bp downstream of a strong pan-neural *nSyb* promoter/enhancer construct. Expression of constructs were validated by IHC against Gba1b, V5, or mCherry, as appropriate.

#### Polyclonal rabbit Gba1b antibody generation

The entire Gba1b protein minus the signal peptide (AA #22-566) was cloned into a GST-vector containing plasmid, induced and purified from inclusion bodies by Abclonal. Three New Zealand rabbits were selected based on low-background expression of pre-immunization bleeds and then immunized with Gba1b with 4 immunizations spanning 40 days. Sera was collected at 80 days and checked by Elisa for antigen reactivity. We screened test-bleeds against Gba1b-overexpressing flies and identified two bleeds which gave strong reactivity against Gba1b-specific overexpressing cells via IHC. One serum was selected for purification by antigen affinity chromatography, which labeled the Gba1b antigen at 89KD by Western Blot. While anti-Gba1b detects overexpressed Gba1b well, endogenous Gba1b expression is extremely low (Davie et al., 2018) and did not emerge above background in *gba1b^Δ1/TT^* or *gba1b^ΔTT/ΔTT^*mutant brains; it is also possible that anti-Gba1b cross-reacts with Gba1a, which is expressed at low levels in Kenyon Cells (Davie et al., 2018).

#### LysoTracker Brain Explant Imaging

Flies in batches of 15 were anesthetized on ice and transferred to an immobilizing plastic collar. Brains were dissected under cold 1X dissection saline (103mM NaCl, 3mM KCL, 5mM TES, 1mM NaH_2_PO_4_, 4mM MgCl) and placed in Terasaki plates containing 1X dissection saline before individual transfer to 12µL of freshly diluted LysoTracker solution (1:500 dilution from stock LysoTracker Red DND-99, 2µM, Thermofisher) for 2 minutes before immediate transfer to 100µL of saline on a microscope slide. Brains were pressed to the bottom of the saline bubble and oriented with dorsal side up (apposed to the coverslip). Brains were immediately imaged on a Leica SP8 confocal using a 40X Lens (N.A. 1.30) at 3X digital zoom. Z-stacks of 5 slices through the cortical region were acquired from optic lobes. Batches of 5 brains (interleaving control and experimental brains) were transferred to individual wells of Saline, LysoTracker, and individual slides in parallel. For co-staining LysoTracker with MagicRed, Nile Red, or in *Lamp^3XmCherry^* (Hegedűs et al., 2016) backgrounds, LysoTracker Green DND-26 was used (also at 2µM, Thermofisher). Here, brains were incubated 2 minutes in LysoTracker green and then 2 minutes in either MagicRed (ab270772 Abcam, reconstituted in 200uL of DMSO) or Nile Red (N1142 Thermo Fisher Scientific, 1mg/mL acetone stock used 1:100 (10µg/mL final) in dissection saline) before mounting as above and immediately imaging. For imaging *myrTdT* and *GFP* flies, we used Lysotracker DeepRed, L12492 (Thermofisher) at 1:200 in dissection saline with native fluorescence for TdTomato and GFP. We were unable to reliably fix LysoTracker signals from *gba1b* brains.

#### IHC Dissections and Staining

As with LysoTracker, flies were anesthetized on ice and immobilized in dissection collars. The proboscis was removed under cold dissection saline and then freshly diluted 4% paraformaldehyde (PFA) was added to the dissection collar (32% EM grade PFA (EMS)). Brains were fixed for 25 minutes; after 5 minutes of immersion in 4% PFA, the remaining head cuticle and surrounding fat was gently removed. Post-fixation, brains were washed three times in 1X PBS before completing the dissection in collars and removing brains into Terasaki wells with 0.5%PBSTx (TritonX-100 0.5% in 1X PBS; for GlcCer and Nile Red staining, 0.05% PBSTx was used to better preserve lipids). Brains were permeabilized for 30 mins then transferred to blocking solution (10%NGS in 0.5%PBSTx) for 40 minutes before adding primary antibodies in 0.5%PBSTx+10%NGS (1:10 CSP, 1:200 anti-FK2 polyubiquitin, 1:500 anti-p62 (Abcam ab178440), 1:20 nc82, 1:50 anti-PDF, 1:2000 anti-Gba1b, 1:10,000 anti-GFP (Abcam ab13970), 1:500 anti-GlcCer (RAS_0010 anti-rabbit Glycobiotech) for 24-48 hours at 4°C while rocking gently. Brains were then washed three times in 0.5%PBSTx before transfer to appropriate secondaries (1:500, Thermo Fisher Scientific) and rocked at 4°C overnight. Brains were washed three times in 0.5%PBSTx before placing in 70% glycerol for clearing, then mounted and imaged in Vectashield. Images were collected using a Leica SP8 confocal microscope equipped with a 40X lens (N.A. 1.30) at 3X digital zoom. Z-stacks of 15 slices through the OL-iCh and MB-ca a were acquired to quantify Ubi aggregate formation. For circadian experiments, IHC experiments were independently repeated three times.

The following antibodies were obtained from the Developmental Studies Hybridoma Bank, created by the NICHD of the NIH and maintained at The University of Iowa, Department of Biology, Iowa City, IA 52242: CSP (6D6) developed by S. Benzer; brp (nc82) developed by E. Bruchner; and PDF (C7) developed by J. Blau.

#### LysoTracker/Ubiquitin particle analysis, Data Plotting

Confocal LIF files were imported into FIJI and analyzed with a custom macro. Maximum intensity projections (MIPS) of slices (equal number between control and experimental) were made and thresholded based on pixel intensity (preserving values 2 standard deviations greater than the mean). Particles were analyzed: for LysoTracker and Ubi puncta, we imposed a circularity requirement of 0.4-1 and size requirement of > 20pixels. For chiasm ubiquitin, we imposed a circularity requirement of 0.1-0.7 and size >100pixels. This FIJI macro worked well for puncta particles but occasionally missed chiasm ubiquitin or erroneously called peripheral cells as chiasm ubiquitin; thus, all analyzed MIPS and FIJI-identified particles were validated by eye post-FIJI pipeline. Particle metrics were exported to excel and graphed and analyzed by ANOVA corrected via Tukey’s test for multiple comparisons in Prism GraphPad or Kruskal-Wallis test for comparisons of nonparametric data (generally Ol-iCh chiasm Ubi).

#### Sleep/Activity (DAM) assay and analysis

We used *Drosophila* Activity Monitors (DAMs, Trikinetics). 1-2 day-old flies (typically males but for select experiments virgin females) were loaded into tubes, and sleep data were recorded until day 10. We used data from day 5 to day 10 after developmental sleep completed. Cessation of activity for 5 mins was used as a proxy for sleep; flies that were inactive for 12h prior to endpoint of the analyzed period were inferred to be dead and omitted from analysis. Data were analyzed in ClockLab as well as with a custom Python library that is available at https://github.com/ClandininLab/sleep_analysis. We confirmed that our python script produced comparable results to both ClockLab and ShinyR Dam (https://karolcichewicz.shinyapps.io/shinyr-dam/) (*Table S5*). Statistical analysis was done by ANOVA between control and mutant cumulative sleep or activity binned on daytime vs nighttime; for data that was not equally variate, Welch’s ANOVA was used. For sleep bout length and number, the Mann-Whitney U test was used.

For testing endogenous circadian rhythms in control, *gba1b^Δ^*, and *glia*>*gba1b^cKO^*, 5 day old flies were shifted into DD (following LD entrainment) and analyzed for days 6-11 after 1 day of adaptation using Clocklab’s FFT and Chi-squared analyses. For Chi-squared, p<0.01, bin =6, and 0-28 time-windows were set. For %Rhythmic, we report data using both FFT amplitude <0.001 as well as Chi-squared power <0.01 (*Table S5*).

For sleep deprivation, a VWR 2500X shaker delivered a randomized 2s pulse every 60s from ZT12 to ZT0.

#### sLNv assay and analysis

We entrained flies in LD (12h-light, 12h-dark), dissected them at dawn (ZT0) and dusk (ZT12), and stained brains for anti-PDF (DSHB C7 1:50) and the neuropil counterstain brp (DSHB nc82 1:20) using secondary antibodies Alexa Fluor 633 (A21126) and Alexa Fluor 488 (A21141) (following protocol in “IHC Dissections and Staining”). Both anti-PDF and *CD8-GFP* genetic labeling of sLNv cells faithfully report sLNv morphology and circadian remodeling (Fernández et al., 2008) (Petsakou et al., 2015). We acquired 1024×1024 resolution confocal stacks of ventral sLNv terminals using a Leica SP8 confocal microscope with a 40X immersion lens (N.A. 1.30) at 2X digital zoom, acquiring image planes 1µm apart. Confocal images were exported to FIJI. Analysis of 3D spread was conducted blind to genotype or ZT using Matlab code kindly provided by Justin Blau (Petsakou et al., 2015). We excluded brains where PdfTri neurons were not fully pruned following metamorphosis, as these PDF+ cells overlap with sLNv terminals (Helfrich-Förster, 1997). Experiments were normalized to highest-volume control condition (typically ZT0 controls) run in parallel with the genetic manipulation, as batch ICH effects changed baseline control volume parameters. All ZT0/ZT12 experiments were dissected on the same day.

#### Lipidomics and Lipid Analyses

Fly cohorts of control (*CS/ISOD1)* and *gba1b^Δ1/TT^*mutants were housed in antiphase incubators and dissected at ZT16-18 (night/asleep) or ZT10-12 (day/awake, evening peak of activity) at day 3 and day 10. For figures were ZT is not indicated, genotypes were pooled across the two dissection ZTs. Crosses were maintained in 12:12 light/dark (LD) incubators that were at 25°C and 50-70% humidity. Light intensity was estimated to be ~400 lumens. For dissections during ‘night’, flies were removed from incubators in aluminum-foil enclosed boxes and not exposed to light until dissection.

15 brains per condition (genotype/timepoint) were dissected in quadruplicate (4 separate tubes). Dissections were in 1X dissection saline (see “IHC dissections” above). Flies were anesthetized on ice; all non-brain tissue (fat/hemocytes, larger trachea) was removed, as well as retina (entirely) and most lamina. Single dissected brains were immediately transferred to Eppendorf tubes containing 20µl saline on ice; after 15 brains were added (10-15mins), 180µl of methanol was added (90% methanol v/v) and brains were snap-frozen on dry ice and stored at −80°C until further analysis. The main lipidomics experiment conducted across day3/day10 was duplicated two months later with independent genetic crosses.

From each group of 15 brains, 10 brains were analyzed for sphingolipids and phospholipids. Brain total lipids containing internal standards (ISTD) were extracted as described by *(61)*. ISTD used included GlucosylßCeramide d18:1/8:0 (Avanti #860540), CerPE d18:1/24:0 (Avanti #860067) and Equisplash™ LIPIDOMIX® Quantitative Mass Spec Internal Standard (Avanti #330731). Lipid extracts were separate by normal phase chromatography (Agilent Zorbax RX-Sil column 3.0 x 100 mm, 1.8 µm particle size) using an Agilent 1260 Infinity LC system (Santa Clara, CA). The total LC run time was 38 min (with 5 min post-run) at a flow rate of 0.3 µL/min. Mobile phase A was composed of isopropanol/hexane/water (58:40:2,v/v) with 5mM ammonium acetate and 0.1% acetic acid. Mobile phase B consisted of isopropanol/hexane/water (50:40:10,v/v) with 5mM ammonium acetate and 0.1% acetic acid. Gradient elution consisted of increase from 34 to 36 min until 100% of mobile phase B, and after 36 min the mobile phase decreased to 0% for 2 more min. All MS measurements were obtained using an Agilent 6430 Triple-Quad LC/MS system (Santa Clara, CA) operating in positive ion mode. Gas-phase ions of various lipid species were obtained using electrospray ionization (ESI). The MS was operated in multiple reaction monitoring (MRM). The collision energies vary from 15 to 45 eV. Data acquisition and analysis was carried out using the Agilent Mass Hunter software package.

For lipid species, total ng/lipid species were calculated per brain per tube as well as relative % of species class. We prioritized ng/brain counts for sphingolipid subspecies but only obtained relative % counts for phospholipid subspecies. We report raw values in *Table 3* but for certain figures report values normalized to the highest ng/brain value per experiment run for that sphingolipid class. For initial bulk and subspecies analyses, we normalized values to highest ng/brain value for individual lipid species across the experimental run; total lipids detected (ng/brain) was unchanged between mutants and controls in both experiments.

For principal component analysis (PCA), lipid species were split into groups based on class (Cer, GlcCer, CerPE, PS, PI, PC, PE), filtered for being common to experimental runs and in the top 99% cumulative fraction, and combined after z-scoring within experiment for individual species (e.g. z-scoring GlcCer 14:1/18:0 across all tubes from experiment #1, then combining with z-scored values for GlcCer 14:1/18:0 from experiment #2). Z-scored matrices were used for PCA analysis in R. To ascertain effects of *gba1b* on lipid species, biplots for PC1-10 from day 10 control and mutant brains (all ZTs) were plotted and colored by genotype. To test for time-effects, control samples from day 10 were analyzed by PCA, and PC1-10 biplots were colored by time. To directly identify time-modulated species, we constructed mean matrices for ‘awake’ and ‘asleep’ z-scores from controls or mutants, and subtracted these matrices to find species that changed most across time. We then calculated t-tests for these species and corrected for multiple comparisons by the Benjamini-Hochberg FDR method. Species with FDR<0.05 were plotted in ggplot2; we confirmed that each identified species showed circadian fluctuations in both raw and normalized data in both experiments.

## Notes

### Competing Interest Statement

The authors have declared no competing interest.

## References

Abruzzi, K.C., Rodriguez, J., Menet, J.S., Desrochers, J., Zadina, A., Luo, W., Tkachev, S., and Rosbash, M. (2011). *Drosophila* CLOCK target gene characterization: implications for circadian tissue-specific gene expression. Genes Dev. 25, 2374–2386. https://doi.org/10.1101/gad.178079.111.

Abruzzi, K.C., Zadina, A., Luo, W., Wiyanto, E., Rahman, R., Guo, F., Shafer, O., and Rosbash, M. (2017). RNA-seq analysis of Drosophila clock and non-clock neurons reveals neuron-specific cycling and novel candidate neuropeptides. PLOS Genetics 13, e1006613. https://doi.org/10.1371/journal.pgen.1006613.

Acharya, U., and Acharya, J.K. (2005). Enzymes of sphingolipid metabolism in Drosophila melanogaster. Cell Mol Life Sci 62, 128–142. https://doi.org/10.1007/s00018-004-4254-1.

Acharya, U., Patel, S., Koundakjian, E., Nagashima, K., Han, X., and Acharya, J.K. (2003). Modulating sphingolipid biosynthetic pathway rescues photoreceptor degeneration. Science 299, 1740–1743. https://doi.org/10.1126/science.1080549.

Artiushin, G., Zhang, S.L., Tricoire, H., and Sehgal, A. (2018). Endocytosis at the Drosophila blood–brain barrier as a function for sleep. ELife 7, e43326. https://doi.org/10.7554/eLife.43326.

Awad, O., Sarkar, C., Panicker, L.M., Miller, D., Zeng, X., Sgambato, J.A., Lipinski, M.M., and Feldman, R.A. (2015). Altered TFEB-mediated lysosomal biogenesis in Gaucher disease iPSC-derived neuronal cells. Hum Mol Genet 24, 5775–5788. https://doi.org/10.1093/hmg/ddv297.

Barth, M., Hirsch, H.V.B., Meinertzhagen, I.A., and Heisenberg, M. (1997). Experience-Dependent Developmental Plasticity in the Optic Lobe of Drosophila melanogaster. J. Neurosci. 17, 1493–1504. https://doi.org/10.1523/JNEUROSCI.17-04-01493.1997.

Bartlett, B.J., Isakson, P., Lewerenz, J., Sanchez, H., Kotzebue, R.W., Cumming, R.C., Harris, G.L., Nezis, I.P., Schubert, D.R., Simonsen, A., et al. (2011). p62, Ref(2)P and ubiquitinated proteins are conserved markers of neuronal aging, aggregate formation and progressive autophagic defects. Autophagy 7, 572–583. https://doi.org/10.4161/auto.7.6.14943.

Becquet, D., Girardet, C., Guillaumond, F., François-Bellan, A.-M., and Bosler, O. (2008). Ultrastructural plasticity in the rat suprachiasmatic nucleus. Possible involvement in clock entrainment. Glia 56, 294–305. https://doi.org/10.1002/glia.20613.

Bedont, J.L., Toda, H., Shi, M., Park, C.H., Quake, C., Stein, C., Kolesnik, A., and Sehgal, A. (2021). Short and long sleeping mutants reveal links between sleep and macroautophagy. Elife 10, e64140. https://doi.org/10.7554/eLife.64140.

Brancaccio, M., Edwards, M.D., Patton, A.P., Smyllie, N.J., Chesham, J.E., Maywood, E.S., and Hastings, M.H. (2019). Cell-autonomous clock of astrocytes drives circadian behavior in mammals. Science 363, 187–192. https://doi.org/10.1126/science.aat4104.

Cangalaya, C., Stoyanov, S., Fischer, K.-D., and Dityatev, A. (2020). Light-induced engagement of microglia to focally remodel synapses in the adult brain. ELife 9, e58435. https://doi.org/10.7554/eLife.58435.

Castro, B.M., Prieto, M., and Silva, L.C. (2014). Ceramide: A simple sphingolipid with unique biophysical properties. Progress in Lipid Research 54, 53–67. https://doi.org/10.1016/j.plipres.2014.01.004.

Chen, Y.-W., Pedersen, J.W., Wandall, H.H., Levery, S.B., Pizette, S., Clausen, H., and Cohen, S.M. (2007). Glycosphingolipids with extended sugar chain have specialized functions in development and behavior of Drosophila. Developmental Biology 306, 736–749. https://doi.org/10.1016/j.ydbio.2007.04.013.

Choi, C., Cao, G., Tanenhaus, A.K., McCarthy, E. v., Jung, M., Schleyer, W., Shang, Y., Rosbash, M., Yin, J.C.P., and Nitabach, M.N. (2012). Autoreceptor Control of Peptide/Neurotransmitter Corelease from PDF Neurons Determines Allocation of Circadian Activity in Drosophila. Cell Reports 2, 332–344. https://doi.org/10.1016/j.celrep.2012.06.021.

Clark, G.T., Yu, Y., Urban, C.A., Fu, G., Wang, C., Zhang, F., Linhardt, R.J., and Hurley, J.M. (2022). Circadian control of heparan sulfate levels times phagocytosis of amyloid beta aggregates. PLOS Genetics 18, e1009994. https://doi.org/10.1371/journal.pgen.1009994.

Dahlgaard, K., Jung, A., Qvortrup, K., Clausen, H., Kjaerulff, O., and Wandall, H.H. (2012). Neurofibromatosis-like phenotype in Drosophila caused by lack of glucosylceramide extension. Proc Natl Acad Sci U S A 109, 6987–6992. https://doi.org/10.1073/pnas.1115453109.

Dasgupta, U., Bamba, T., Chiantia, S., Karim, P., Tayoun, A.N.A., Yonamine, I., Rawat, S.S., Rao, R.P., Nagashima, K., Fukusaki, E., et al. (2009). Ceramide kinase regulates phospholipase C and phosphatidylinositol 4, 5, bisphosphate in phototransduction. PNAS 106, 20063–20068. https://doi.org/10.1073/pnas.0911028106.

Davie, K., Janssens, J., Koldere, D., Waegeneer, M.D., Pech, U., Kreft, Ł., Aibar, S., Makhzami, S., Christiaens, V., González-Blas, C.B., et al. (2018). A Single-Cell Transcriptome Atlas of the Aging Drosophila Brain. Cell 174, 982–998.e20. https://doi.org/10.1016/j.cell.2018.05.057.

Davis, M.Y., Trinh, K., Thomas, R.E., Yu, S., Germanos, A.A., Whitley, B.N., Sardi, S.P., Montine, T.J., and Pallanck, L.J. (2016). Glucocerebrosidase Deficiency in Drosophila Results in α-Synuclein-Independent Protein Aggregation and Neurodegeneration. PLOS Genetics 12, e1005944. https://doi.org/10.1371/journal.pgen.1005944.

Doherty, J., Logan, M.A., Tasdemir, O.E., and Freeman, M.R. (2009). Ensheathing Glia Function as Phagocytes in the Adult Drosophila Brain. Journal of Neuroscience 29, 4768–4781. https://doi.org/10.1523/JNEUROSCI.5951-08.2009.

Enquist, I.B., Lo Bianco, C., Ooka, A., Nilsson, E., Månsson, J.-E., Ehinger, M., Richter, J., Brady, R.O., Kirik, D., and Karlsson, S. (2007). Murine models of acute neuronopathic Gaucher disease. Proc Natl Acad Sci U S A 104, 17483–17488. https://doi.org/10.1073/pnas.0708086104.

Fernández, M.P., Berni, J., and Ceriani, M.F. (2008). Circadian remodeling of neuronal circuits involved in rhythmic behavior. PLoS Biol 6, e69. https://doi.org/10.1371/journal.pbio.0060069.

Fitzner, D., Bader, J.M., Penkert, H., Bergner, C.G., Su, M., Weil, M.-T., Surma, M.A., Mann, M., Klose, C., and Simons, M. (2020). Cell-Type- and Brain-Region-Resolved Mouse Brain Lipidome. Cell Reports 32, 108132. https://doi.org/10.1016/j.celrep.2020.108132.

Freeman, M.R. (2015). Drosophila Central Nervous System Glia. Cold Spring Harb Perspect Biol 7, a020552. https://doi.org/10.1101/cshperspect.a020552.

Futerman, A.H., and van Meer, G. (2004). The cell biology of lysosomal storage disorders. Nat Rev Mol Cell Biol 5, 554–565. https://doi.org/10.1038/nrm1423.

Gan-Or, Z., Mirelman, A., Postuma, R.B., Arnulf, I., Bar-Shira, A., Dauvilliers, Y., Desautels, A., Gagnon, J.-F., Leblond, C.S., Frauscher, B., et al. (2015). GBA mutations are associated with Rapid Eye Movement Sleep Behavior Disorder. Ann Clin Transl Neurol 2, 941–945. https://doi.org/10.1002/acn3.228.

Ghosh, A., Kling, T., Snaidero, N., Sampaio, J.L., Shevchenko, A., Gras, H., Geurten, B., Göpfert, M.C., Schulz, J.B., Voigt, A., et al. (2013). A Global In Vivo Drosophila RNAi Screen Identifies a Key Role of Ceramide Phosphoethanolamine for Glial Ensheathment of Axons. PLOS Genetics 9, e1003980. https://doi.org/10.1371/journal.pgen.1003980.

Grima, B., Chélot, E., Xia, R., and Rouyer, F. (2004). Morning and evening peaks of activity rely on different clock neurons of the Drosophila brain. Nature 431, 869–873. https://doi.org/10.1038/nature02935.

Guan, X.L., Cestra, G., Shui, G., Kuhrs, A., Schittenhelm, R.B., Hafen, E., van der Goot, F.G., Robinett, C.C., Gatti, M., Gonzalez-Gaitan, M., et al. (2013). Biochemical membrane lipidomics during Drosophila development. Dev Cell 24, 98–111. https://doi.org/10.1016/j.devcel.2012.11.012.

Guttenplan, K.A., Weigel, M.K., Prakash, P., Wijewardhane, P.R., Hasel, P., Rufen-Blanchette, U., Münch, A.E., Blum, J.A., Fine, J., Neal, M.C., et al. (2021). Neurotoxic reactive astrocytes induce cell death via saturated lipids. Nature 599, 102–107. https://doi.org/10.1038/s41586-021-03960-y.

Haines, N., and Irvine, K.D. (2005). Functional analysis of Drosophila beta1,4-N-acetlygalactosaminyltransferases. Glycobiology 15, 335–346. https://doi.org/10.1093/glycob/cwi017.

Hegedűs, K., Takáts, S., Boda, A., Jipa, A., Nagy, P., Varga, K., Kovács, A.L., and Juhász, G. (2016). The Ccz1-Mon1-Rab7 module and Rab5 control distinct steps of autophagy. MBoC 27, 3132–3142. https://doi.org/10.1091/mbc.e16-03-0205.

Heisenberg, M., Heusipp, M., and Wanke, C. (1995). Structural plasticity in the Drosophila brain. J. Neurosci. 15, 1951–1960. https://doi.org/10.1523/JNEUROSCI.15-03-01951.1995.

Helfrich-Förster, C. (1997). Development of pigment-dispersing hormone-immunoreactive neurons in the nervous system of Drosophila melanogaster. Journal of Comparative Neurology 380, 335–354. https://doi.org/10.1002/(SICI)1096-9861(19970414)380:3<335::AID-CNE4>3.0.CO;2-3.

Helfrich-Förster, C. (2000). Differential Control of Morning and Evening Components in the Activity Rhythm of Drosophila melanogaster—Sex-Specific Differences Suggest a Different Quality of Activity. J Biol Rhythms 15, 135–154. https://doi.org/10.1177/074873040001500208.

Herrero, A., Duhart, J.M., and Ceriani, M.F. (2017). Neuronal and Glial Clocks Underlying Structural Remodeling of Pacemaker Neurons in Drosophila. Front Physiol 8, 918. https://doi.org/10.3389/fphys.2017.00918.

Huang, Y., Huang, S., Lam, S.M., Liu, Z., Shui, G., and Zhang, Y.Q. (2016). Acsl, the Drosophila ortholog of intellectual-disability-related ACSL4, inhibits synaptic growth by altered lipids. J Cell Sci 129, 4034–4045. https://doi.org/10.1242/jcs.195032.

Huang, Y., Huang, S., Di Scala, C., Wang, Q., Wandall, H.H., Fantini, J., and Zhang, Y.Q. (2018). The glycosphingolipid MacCer promotes synaptic bouton formation in Drosophila by interacting with Wnt. ELife 7, e38183. https://doi.org/10.7554/eLife.38183.

Jewett, K.A., Thomas, R.E., Phan, C.Q., Lin, B., Milstein, G., Yu, S., Bettcher, L.F., Neto, F.C., Djukovic, D., Raftery, D., et al. (2021). Glucocerebrosidase reduces the spread of protein aggregation in a Drosophila melanogaster model of neurodegeneration by regulating proteins trafficked by extracellular vesicles. PLoS Genet 17, e1008859. https://doi.org/10.1371/journal.pgen.1008859.

Jung, W., Liu, C., Yu, Y., Chang, Y., Lien, W., Chao, H., Huang, S., Kuo, C., Ho, H., and Chan, C. (2017). Lipophagy prevents activity-dependent neurodegeneration due to dihydroceramide accumulation in vivo. EMBO Rep 18, 1150–1165. https://doi.org/10.15252/embr.201643480.

Katewa, S.D., Akagi, K., Bose, N., Rakshit, K., Camarella, T., Zheng, X., Hall, D., Davis, S., Nelson, C.S., Brem, R.B., et al. (2016). Peripheral circadian clocks mediate dietary restriction dependent changes in lifespan and fat metabolism in Drosophila. Cell Metab 23, 143–154. https://doi.org/10.1016/j.cmet.2015.10.014.

Kawasaki, H., Suzuki, T., Ito, K., Takahara, T., Goto-Inoue, N., Setou, M., Sakata, K., and Ishida, N. (2017). Minos-insertion mutant of the Drosophila GBA gene homologue showed abnormal phenotypes of climbing ability, sleep and life span with accumulation of hydroxy-glucocerebroside. Gene 614, 49–55. https://doi.org/10.1016/j.gene.2017.03.004.

Keatinge, M., Bui, H., Menke, A., Chen, Y.-C., Sokol, A.M., Bai, Q., Ellett, F., Da Costa, M., Burke, D., Gegg, M., et al. (2015). Glucocerebrosidase 1 deficient Danio rerio mirror key pathological aspects of human Gaucher disease and provide evidence of early microglial activation preceding alpha-synuclein-independent neuronal cell death. Hum Mol Genet 24, 6640–6652. https://doi.org/10.1093/hmg/ddv369.

Kinghorn, K.J., Grönke, S., Castillo-Quan, J.I., Woodling, N.S., Li, L., Sirka, E., Gegg, M., Mills, K., Hardy, J., Bjedov, I., et al. (2016). A Drosophila Model of Neuronopathic Gaucher Disease Demonstrates Lysosomal-Autophagic Defects and Altered mTOR Signalling and Is Functionally Rescued by Rapamycin. J Neurosci 36, 11654–11670. https://doi.org/10.1523/JNEUROSCI.4527-15.2016.

Koh, K., Joiner, W.J., Wu, M.N., Yue, Z., Smith, C.J., and Sehgal, A. (2008). Identification of SLEEPLESS, a Sleep-Promoting Factor. Science 321, 372–376. https://doi.org/10.1126/science.1155942.

Konopka, R.J., and Benzer, S. (1971). Clock Mutants of Drosophila melanogaster. PNAS 68, 2112–2116. https://doi.org/10.1073/pnas.68.9.2112.

Kozlov, A., Koch, R., and Nagoshi, E. (2020). Nitric oxide mediates neuro-glial interaction that shapes Drosophila circadian behavior. PLOS Genetics 16, e1008312. https://doi.org/10.1371/journal.pgen.1008312.

Kremer, M.C., Jung, C., Batelli, S., Rubin, G.M., and Gaul, U. (2017). The glia of the adult *Drosophila* nervous system: Glia Anatomy in Adult Drosophila Nervous System. Glia 65, 606–638. https://doi.org/10.1002/glia.23115.

Krohn, L., Ruskey, J.A., Rudakou, U., Leveille, E., Asayesh, F., Hu, M.T.M., Arnulf, I., Dauvilliers, Y., Högl, B., Stefani, A., et al. (2020). GBA variants in REM sleep behavior disorder: A multicenter study. Neurology 95, e1008–e1016. https://doi.org/10.1212/WNL.0000000000010042.

Krzeptowski, W., Hess, G., and Pyza, E. (2018). Circadian Plasticity in the Brain of Insects and Rodents. Frontiers in Neural Circuits 12, 32. https://doi.org/10.3389/fncir.2018.00032.

Kunduri, G., Turner-Evans, D., Konya, Y., Izumi, Y., Nagashima, K., Lockett, S., Holthuis, J., Bamba, T., Acharya, U., and Acharya, J.K. (2018). Defective cortex glia plasma membrane structure underlies light-induced epilepsy in cpes mutants. Proc Natl Acad Sci U S A 115, E8919–E8928. https://doi.org/10.1073/pnas.1808463115.

Kurmangaliyev, Y.Z., Yoo, J., Valdes-Aleman, J., Sanfilippo, P., and Zipursky, S.L. (2020). Transcriptional Programs of Circuit Assembly in the Drosophila Visual System. Neuron 108, 1045–1057.e6. https://doi.org/10.1016/j.neuron.2020.10.006.

Lee, P.-T., Zirin, J., Kanca, O., Lin, W.-W., Schulze, K.L., Li-Kroeger, D., Tao, R., Devereaux, C., Hu, Y., Chung, V., et al. (2018). A gene-specific T2A-GAL4 library for Drosophila. ELife 7, e35574. https://doi.org/10.7554/eLife.35574.

Leng, Y., Musiek, E.S., Hu, K., Cappuccio, F.P., and Yaffe, K. (2019). Association between circadian rhythms and neurodegenerative diseases. Lancet Neurol 18, 307–318. https://doi.org/10.1016/S1474-4422(18)30461-7.

Liddelow, S.A., Guttenplan, K.A., Clarke, L.E., Bennett, F.C., Bohlen, C.J., Schirmer, L., Bennett, M.L., Münch, A.E., Chung, W.-S., Peterson, T.C., et al. (2017). Neurotoxic reactive astrocytes are induced by activated microglia. Nature 541, 481–487. https://doi.org/10.1038/nature21029.

Lin, G., Lee, P.-T., Chen, K., Mao, D., Tan, K.L., Zuo, Z., Lin, W.-W., Wang, L., and Bellen, H.J. (2018). Phospholipase PLA2G6, a Parkinsonism-Associated Gene, Affects Vps26 and Vps35, Retromer Function, and Ceramide Levels, Similar to α-Synuclein Gain. Cell Metabolism 28, 605–618.e6. https://doi.org/10.1016/j.cmet.2018.05.019.

Lin, G., Wang, L., Marcogliese, P.C., and Bellen, H.J. (2019). Sphingolipids in the Pathogenesis of Parkinson’s Disease and Parkinsonism. Trends in Endocrinology & Metabolism 30, 106–117. https://doi.org/10.1016/j.tem.2018.11.003.

Liu, G., Boot, B., Locascio, J.J., Jansen, I.E., Winder-Rhodes, S., Eberly, S., Elbaz, A., Brice, A., Ravina, B., van Hilten, J.J., et al. (2016). Specifically neuropathic Gaucher’s mutations accelerate cognitive decline in Parkinson’s. Ann Neurol 80, 674–685. https://doi.org/10.1002/ana.24781.

Liu, H., Wang, X., Chen, L., Chen, L., Tsirka, S.E., Ge, S., and Xiong, Q. (2021). Microglia modulate stable wakefulness via the thalamic reticular nucleus in mice. Nat Commun 12, 4646. https://doi.org/10.1038/s41467-021-24915-x.

Liu, L., Zhang, K., Sandoval, H., Yamamoto, S., Jaiswal, M., Sanz, E., Li, Z., Hui, J., Graham, B.H., Quintana, A., et al. (2015). Glial lipid droplets and ROS induced by mitochondrial defects promote neurodegeneration. Cell 160, 177–190. https://doi.org/10.1016/j.cell.2014.12.019.

Mazzulli, J.R., Xu, Y.-H., Sun, Y., Knight, A.L., McLean, P.J., Caldwell, G.A., Sidransky, E., Grabowski, G.A., and Krainc, D. (2011). Gaucher Disease Glucocerebrosidase and α-Synuclein Form a Bidirectional Pathogenic Loop in Synucleinopathies. Cell 146, 37–52. https://doi.org/10.1016/j.cell.2011.06.001.

Meivar-Levy, I., Sabanay, H., Bershadsky, A.D., and Futerman, A.H. (1997). The Role of Sphingolipids in the Maintenance of Fibroblast Morphology: THE INHIBITION OF PROTRUSIONAL ACTIVITY, CELL SPREADING, AND CYTOKINESIS INDUCED BY FUMONISIN B1 CAN BE REVERSED BY GANGLIOSIDE GM3*. Journal of Biological Chemistry 272, 1558–1564. https://doi.org/10.1074/jbc.272.3.1558.

Merrill, A.H. (2011). Sphingolipid and Glycosphingolipid Metabolic Pathways in the Era of Sphingolipidomics. Chem Rev 111, 6387–6422. https://doi.org/10.1021/cr2002917.

Mikulka, C.R., Dearborn, J.T., Benitez, B.A., Strickland, A., Liu, L., Milbrandt, J., and Sands, M.S. (2020). Cell-autonomous expression of the acid hydrolase galactocerebrosidase. Proc Natl Acad Sci USA 117, 9032–9041. https://doi.org/10.1073/pnas.1917675117.

Musashe, D.T., Purice, M.D., Speese, S.D., Doherty, J., and Logan, M.A. (2016). Insulin-like Signaling Promotes Glial Phagocytic Clearance of Degenerating Axons through Regulation of Draper. Cell Reports 16, 1838–1850. https://doi.org/10.1016/j.celrep.2016.07.022.

Ng, F.S., and Jackson, F.R. (2015). The ROP vesicle release factor is required in adult Drosophila glia for normal circadian behavior. Front Cell Neurosci 9, 256. https://doi.org/10.3389/fncel.2015.00256.

O’Brien, J.S., and Sampson, E.L. (1965). Lipid composition of the normal human brain: gray matter, white matter, and myelin. J Lipid Res 6, 537–544.

Osellame, L.D., Rahim, A.A., Hargreaves, I.P., Gegg, M.E., Richard-Londt, A., Brandner, S., Waddington, S.N., Schapira, A.H.V., and Duchen, M.R. (2013). Mitochondria and Quality Control Defects in a Mouse Model of Gaucher Disease—Links to Parkinson’s Disease. Cell Metab 17, 941–953. https://doi.org/10.1016/j.cmet.2013.04.014.

Pan, T., Kondo, S., Le, W., and Jankovic, J. (2008). The role of autophagy-lysosome pathway in neurodegeneration associated with Parkinson’s disease. Brain 131, 1969–1978. https://doi.org/10.1093/brain/awm318.

Parisky, K.M., Agosto, J., Pulver, S.R., Shang, Y., Kuklin, E., Hodge, J.J.L., Kang, K., Liu, X., Garrity, P.A., Rosbash, M., et al. (2008). PDF Cells Are a GABA-Responsive Wake-Promoting Component of the Drosophila Sleep Circuit. Neuron 60, 672–682. https://doi.org/10.1016/j.neuron.2008.10.042.

Park, J.H., Helfrich-Förster, C., Lee, G., Liu, L., Rosbash, M., and Hall, J.C. (2000). Differential regulation of circadian pacemaker output by separate clock genes in Drosophila. PNAS 97, 3608–3613. https://doi.org/10.1073/pnas.97.7.3608.

Petsakou, A., Sapsis, T.P., and Blau, J. (2015). Circadian Rhythms in Rho1 Activity Regulate Neuronal Plasticity and Network Hierarchy. Cell 162, 823–835. https://doi.org/10.1016/j.cell.2015.07.010.

Pfeiffer, B.D., Jenett, A., Hammonds, A.S., Ngo, T.-T.B., Misra, S., Murphy, C., Scully, A., Carlson, J.W., Wan, K.H., Laverty, T.R., et al. (2008). Tools for neuroanatomy and neurogenetics in Drosophila. PNAS 105, 9715–9720. https://doi.org/10.1073/pnas.0803697105.

Pircs, K., Nagy, P., Varga, A., Venkei, Z., Erdi, B., Hegedus, K., and Juhasz, G. (2012). Advantages and Limitations of Different p62-Based Assays for Estimating Autophagic Activity in Drosophila. PLOS ONE 7, e44214. https://doi.org/10.1371/journal.pone.0044214.

Pyza, E., and Meinertzhagen, I.A. (1999). Daily rhythmic changes of cell size and shape in the first optic neuropil in Drosophila melanogaster. Journal of Neurobiology 40, 77–88. https://doi.org/10.1002/(SICI)1097-4695(199907)40:1<77::AID-NEU7>3.0.CO;2-0.

Raju, D., Schonauer, S., Hamzeh, H., Flynn, K.C., Bradke, F., Dorp, K. vom, Dörmann, P., Yildiz, Y., Trötschel, C., Poetsch, A., et al. (2015). Accumulation of Glucosylceramide in the Absence of the Beta-Glucosidase GBA2 Alters Cytoskeletal Dynamics. PLOS Genetics 11, e1005063. https://doi.org/10.1371/journal.pgen.1005063.

Ross, C.A., and Poirier, M.A. (2004). Protein aggregation and neurodegenerative disease. Nat Med 10, S10–S17. https://doi.org/10.1038/nm1066.

Schafer, D.P., Lehrman, E.K., Kautzman, A.G., Koyama, R., Mardinly, A.R., Yamasaki, R., Ransohoff, R.M., Greenberg, M.E., Barres, B.A., and Stevens, B. (2012). Microglia Sculpt Postnatal Neural Circuits in an Activity and Complement-Dependent Manner. Neuron 74, 691–705. https://doi.org/10.1016/j.neuron.2012.03.026.

Schöndorf, D.C., Aureli, M., McAllister, F.E., Hindley, C.J., Mayer, F., Schmid, B., Sardi, S.P., Valsecchi, M., Hoffmann, S., Schwarz, L.K., et al. (2014). iPSC-derived neurons from GBA1-associated Parkinson’s disease patients show autophagic defects and impaired calcium homeostasis. Nat Commun 5, 4028. https://doi.org/10.1038/ncomms5028.

Sidransky, E., and Lopez, G. (2012). The link between the GBA gene and parkinsonism. Lancet Neurol 11, 986–998. https://doi.org/10.1016/S1474-4422(12)70190-4.

Sidransky, E., Nalls, M.A., Aasly, J.O., Aharon-Peretz, J., Annesi, G., Barbosa, E.R., Bar-Shira, A., Berg, D., Bras, J., Brice, A., et al. (2009). Multicenter analysis of glucocerebrosidase mutations in Parkinson’s disease. N Engl J Med 361, 1651–1661. https://doi.org/10.1056/NEJMoa0901281.

Sivachenko, A., Li, Y., Abruzzi, K.C., and Rosbash, M. (2013). The transcription factor Mef2 links the Drosophila core clock to Fas2, neuronal morphology, and circadian behavior. Neuron 79, 281–292. https://doi.org/10.1016/j.neuron.2013.05.015.

Soller, M., Haussmann, I.U., Hollmann, M., Choffat, Y., White, K., Kubli, E., and Schäfer, M.A. (2006). Sex-Peptide-Regulated Female Sexual Behavior Requires a Subset of Ascending Ventral Nerve Cord Neurons. Current Biology 16, 1771–1782. https://doi.org/10.1016/j.cub.2006.07.055.

Stanhope, B.A., Jaggard, J.B., Gratton, M., Brown, E.B., and Keene, A.C. (2020). Sleep Regulates Glial Plasticity and Expression of the Engulfment Receptor Draper Following Neural Injury. Current Biology 30, 1092–1101.e3. https://doi.org/10.1016/j.cub.2020.02.057.

Stavropoulos, N., and Young, M.W. (2011). insomniac and Cullin-3 Regulate Sleep and Wakefulness in Drosophila. Neuron 72, 964–976. https://doi.org/10.1016/j.neuron.2011.12.003.

Suresh, S.N., Verma, V., Sateesh, S., Clement, J.P., and Manjithaya, R. (2018). Neurodegenerative diseases: model organisms, pathology and autophagy. J Genet 97, 679–701. https://doi.org/10.1007/s12041-018-0955-3.

Tainton-Heap, L.A.L., Kirszenblat, L.C., Notaras, E.T., Grabowska, M.J., Jeans, R., Feng, K., Shaw, P.J., and van Swinderen, B. (2021). A Paradoxical Kind of Sleep in Drosophila melanogaster. Current Biology 31, 578–590.e6. https://doi.org/10.1016/j.cub.2020.10.081.

Tsai, J.W., Kostyleva, R., Chen, P.-L., Rivas-Serna, I.M., Clandinin, M.T., Meinertzhagen, I.A., and Clandinin, T.R. (2019). Transcriptional Feedback Links Lipid Synthesis to Synaptic Vesicle Pools in Drosophila Photoreceptors. Neuron 101, 721–737.e4. https://doi.org/10.1016/j.neuron.2019.01.015.

Uemura, N., Koike, M., Ansai, S., Kinoshita, M., Ishikawa-Fujiwara, T., Matsui, H., Naruse, K., Sakamoto, N., Uchiyama, Y., Todo, T., et al. (2015). Viable Neuronopathic Gaucher Disease Model in Medaka (Oryzias latipes) Displays Axonal Accumulation of Alpha-Synuclein. PLoS Genet 11. https://doi.org/10.1371/journal.pgen.1005065.

Valadas, J.S., Esposito, G., Vandekerkhove, D., Miskiewicz, K., Deaulmerie, L., Raitano, S., Seibler, P., Klein, C., and Verstreken, P. (2018). ER Lipid Defects in Neuropeptidergic Neurons Impair Sleep Patterns in Parkinson’s Disease. Neuron 98, 1155–1169.e6. https://doi.org/10.1016/j.neuron.2018.05.022.

Woeste, M.A., Stern, S., Raju, D.N., Grahn, E., Dittmann, D., Gutbrod, K., Dörmann, P., Hansen, J.N., Schonauer, S., Marx, C.E., et al. (2019). Species-specific differences in nonlysosomal glucosylceramidase GBA2 function underlie locomotor dysfunction arising from loss-of-function mutations. J Biol Chem 294, 3853–3871. https://doi.org/10.1074/jbc.RA118.006311.

Xie, L., Kang, H., Xu, Q., Chen, M.J., Liao, Y., Thiyagarajan, M., O’Donnell, J., Christensen, D.J., Nicholson, C., Iliff, J.J., et al. (2013). Sleep Drives Metabolite Clearance from the Adult Brain. Science 342, 10.1126/science.1241224. https://doi.org/10.1126/science.1241224.

Yadav, R.S., and Tiwari, N.K. (2014). Lipid Integration in Neurodegeneration: An Overview of Alzheimer’s Disease. Mol Neurobiol 50, 168–176. https://doi.org/10.1007/s12035-014-8661-5.

Yildirim, K., Petri, J., Kottmeier, R., and Klämbt, C. (2019). Drosophila glia: Few cell types and many conserved functions. Glia 67, 5–26. https://doi.org/10.1002/glia.23459.

Yin, J., Gibbs, M., Long, C., Rosenthal, J., Kim, H.S., Kim, A., Sheng, C., Ding, P., Javed, U., and Yuan, Q. (2018). Transcriptional Regulation of Lipophorin Receptors Supports Neuronal Adaptation to Chronic Elevations of Activity. Cell Reports 25, 1181–1192.e4. https://doi.org/10.1016/j.celrep.2018.10.016.

Zhang, S.L., Yue, Z., Arnold, D.M., Artiushin, G., and Sehgal, A. (2018). A Circadian Clock in the Blood-Brain Barrier Regulates Xenobiotic Efflux. Cell 173, 130–139.e10. https://doi.org/10.1016/j.cell.2018.02.017.

Zhang, Y., Chen, K., Sloan, S.A., Bennett, M.L., Scholze, A.R., O’Keeffe, S., Phatnani, H.P., Guarnieri, P., Caneda, C., Ruderisch, N., et al. (2014). An RNA-Sequencing Transcriptome and Splicing Database of Glia, Neurons, and Vascular Cells of the Cerebral Cortex. J. Neurosci. 34, 11929–11947. https://doi.org/10.1523/JNEUROSCI.1860-14.2014.

Zhang, Y., Sloan, S.A., Clarke, L.E., Caneda, C., Plaza, C.A., Blumenthal, P.D., Vogel, H., Steinberg, G.K., Edwards, M.S.B., Li, G., et al. (2016). Purification and Characterization of Progenitor and Mature Human Astrocytes Reveals Transcriptional and Functional Differences with Mouse. Neuron 89, 37–53. https://doi.org/10.1016/j.neuron.2015.11.013.

Zuchero, J.B., and Barres, B.A. (2015). Glia in mammalian development and disease. Development 142, 3805–3809. https://doi.org/10.1242/dev.129304.

